# Cereulide synthetase acquisition and loss events within the evolutionary history of Group III *Bacillus cereus sensu lato* facilitate the transition between emetic and diarrheal foodborne pathogen

**DOI:** 10.1101/2020.05.12.090951

**Authors:** Laura M. Carroll, Martin Wiedmann

## Abstract

Cereulide-producing members of *Bacillus cereus sensu lato* (*B. cereus s.l.*) Group III, also known as “emetic *B. cereus*”, possess cereulide synthetase, a plasmid-encoded, non-ribosomal peptide synthetase encoded by the *ces* gene cluster. Despite the documented risks that cereulide-producing strains pose to public health, the level of genomic diversity encompassed by “emetic *B. cereus*” has never been evaluated at a whole-genome scale. Here, we employ a phylogenomic approach to characterize Group III *B. cereus s.l.* genomes which possess *ces* (*ces*-positive) alongside their closely related *ces*-negative counterparts to (i) assess the genomic diversity encompassed by “emetic *B. cereus*”, and (ii) identify potential *ces* loss and/or gain events within the evolutionary history of the high-risk and medically relevant sequence type (ST) 26 lineage often associated with emetic foodborne illness. Using all publicly available *ces*-positive Group III *B. cereus s.l.* genomes and the *ces*-negative genomes interspersed among them (*n* = 150), we show that “emetic *B. cereus*” is not clonal; rather, multiple lineages within Group III harbor cereulide-producing strains, all of which share a common ancestor incapable of producing cereulide (posterior probability [PP] 0.86-0.89). The ST 26 common ancestor was predicted to have emerged as *ces*-negative (PP 0.60-0.93) circa 1904 (95% highest posterior density [HPD] interval 1837.1-1957.8) and first acquired the ability to produce cereulide before 1931 (95% HPD 1893.2-1959.0). Three subsequent *ces* loss events within ST 26 were observed, including among isolates responsible for *B. cereus s.l.* toxicoinfection (i.e., “diarrheal” illness).

**Importance:** “*B. cereus*” is responsible for thousands of cases of foodborne disease each year worldwide, causing two distinct forms of illness: (i) intoxication via cereulide (i.e., “emetic” syndrome) or (ii) toxicoinfection via multiple enterotoxins (i.e., “diarrheal” syndrome). Here, we show that “emetic *B. cereus*” is not a clonal, homogenous unit that resulted from a single cereulide synthetase gain event followed by subsequent proliferation; rather, cereulide synthetase acquisition and loss is a dynamic, ongoing process that occurs across lineages, allowing some Group III *B. cereus s.l.* populations to oscillate between diarrheal and emetic foodborne pathogen over the course of their evolutionary histories. We also highlight the care that must be taken when selecting a reference genome for whole-genome sequencing-based investigation of emetic *B. cereus s.l.* outbreaks, as some reference genome selections can lead to a confounding loss of resolution and potentially hinder epidemiological investigations.

## Introduction

The *Bacillus cereus* group (also known as *B. cereus sensu lato* [*s.l.*]) is a complex of closely related, Gram-positive, spore-forming members of the genus *Bacillus*, which vary in their ability to cause illness in humans (1). Members of *B. cereus s.l.* were estimated to be responsible for more than 256,000 foodborne intoxications worldwide in 2010 (2), although this is likely an underestimate due to the mild symptoms frequently associated with this illness (1). Foodborne “*B. cereus*” intoxication (i.e., “emetic” illness) is caused by cereulide, a highly heat- and pH-stable toxin, which is pre-formed in a food matrix prior to consumption. These intoxications have a relatively short incubation period (typically 0.5 – 6 h) and are often accompanied by symptoms of vomiting and nausea (1, 3–5). This can be contrasted with “*B. cereus*” toxicoinfection (i.e., “diarrheal” illness), a different form of illness in which multiple enterotoxins produced within the host small intestine yield diarrheal symptoms which typically onset after 8 – 16 h (1, 6). Notably, emetic and diarrheal symptoms are not always congruent with “*B. cereus*” emetic and diarrheal syndromes, respectively, as both vomiting and diarrheal symptoms may be reported among cases (7, 8).

Production of cereulide, the toxin responsible for emetic “*B. cereus*” foodborne illness, can be attributed to cereulide synthetase, a non-ribosomal peptide synthetase encoded by the cereulide synthetase biosynthetic gene cluster (*ces*) (9, 10). *ces* has been detected in two major *B. cereus s.l.* phylogenetic groups (assigned using the sequence of pantoate-beta-alanine ligase [*panC*] and a seven-group typing scheme): Group III and Group VI of *B. cereus s.l.* (10–16). While cereulide-producing Group VI strains, also known as “emetic *B. weihenstephanensis*”, have been isolated on rare occasions (14, 15, 17–19), the bulk of cereulide-producing strains belong to Group III (8, 10, 13, 16). Often referred to as “emetic *B. cereus*", cereulide-producing Group III strains often harbor *ces* on plasmids (9, 10, 19), and have been linked to outbreaks around the world (5, 7, 8, 20). It is essential to note that Group III *B. cereus s.l.* isolates do not belong to the *B. cereus sensu stricto* (*s.s.*) species (i.e., *B. cereus s.l.* Group IV) (7, 21). A recently proposed taxonomic reorganization of *B. cereus s.l.* (21) refers to Group III *B. cereus s.l.* as *B. mosaicus*; however, the use of “Group III *B. cereus s.l.*” throughout the remainder of this study is intentional, as, at the present time, it is likely more interpretable to microbiologists than the recently proposed nomenclature.

Despite the documented risks that cereulide-producing strains pose to public health, the level of genomic diversity encompassed by “emetic *B. cereus*” has not been evaluated at a whole-genome scale. Furthermore, potential heterogeneity in cereulide production capabilities among lineages encompassed by “emetic *B. cereus*” has never been assessed; plasmid-encoded *ces* and, thus, the ability to produce cereulide, can hypothetically be gained or lost within a lineage, although the extent to which this happens is unknown. Here, we employ phylogenomic approaches to characterize Group III *B. cereus s.l.* genomes that possess *ces* (*ces*-positive) alongside their closely related *ces*-negative counterparts to (i) assess the genomic diversity encompassed by cereulide-producing Group III strains (i.e., “emetic *B. cereus*”), and (ii) identify potential *ces* loss and/or gain events within the “emetic *B. cereus*” evolutionary history.

## Results

### Cereulide-producing members of Group III *B. cereus s.l.* are distributed across multiple lineages and share a common ancestor incapable of synthesizing cereulide

Of the 2,261 *B. cereus s.l.* genomes queried here (see Supplemental Table S1), 60 genomes belonged to *panC* Group III and possessed cereulide synthetase-encoding *cesABCD* (referred to hereafter as “*ces*-positive” genomes). Overall, 31 STs assigned using *in silico* multi-locus sequence typing (MLST) were observed among the 150 Group III isolates included in this study, with *ces*-positive isolates represented by five STs (ST 26, 144, 164, 869, and 2056; Figure 1 and Supplemental Table S1). Four of these STs (ST 26, 144, 164, and 869) also encompassed one or more isolates that lacked cereulide synthetase (referred to hereafter as “*ces*-negative isolates”; Figure 1 and Supplemental Table S1).

**Figure 1.**
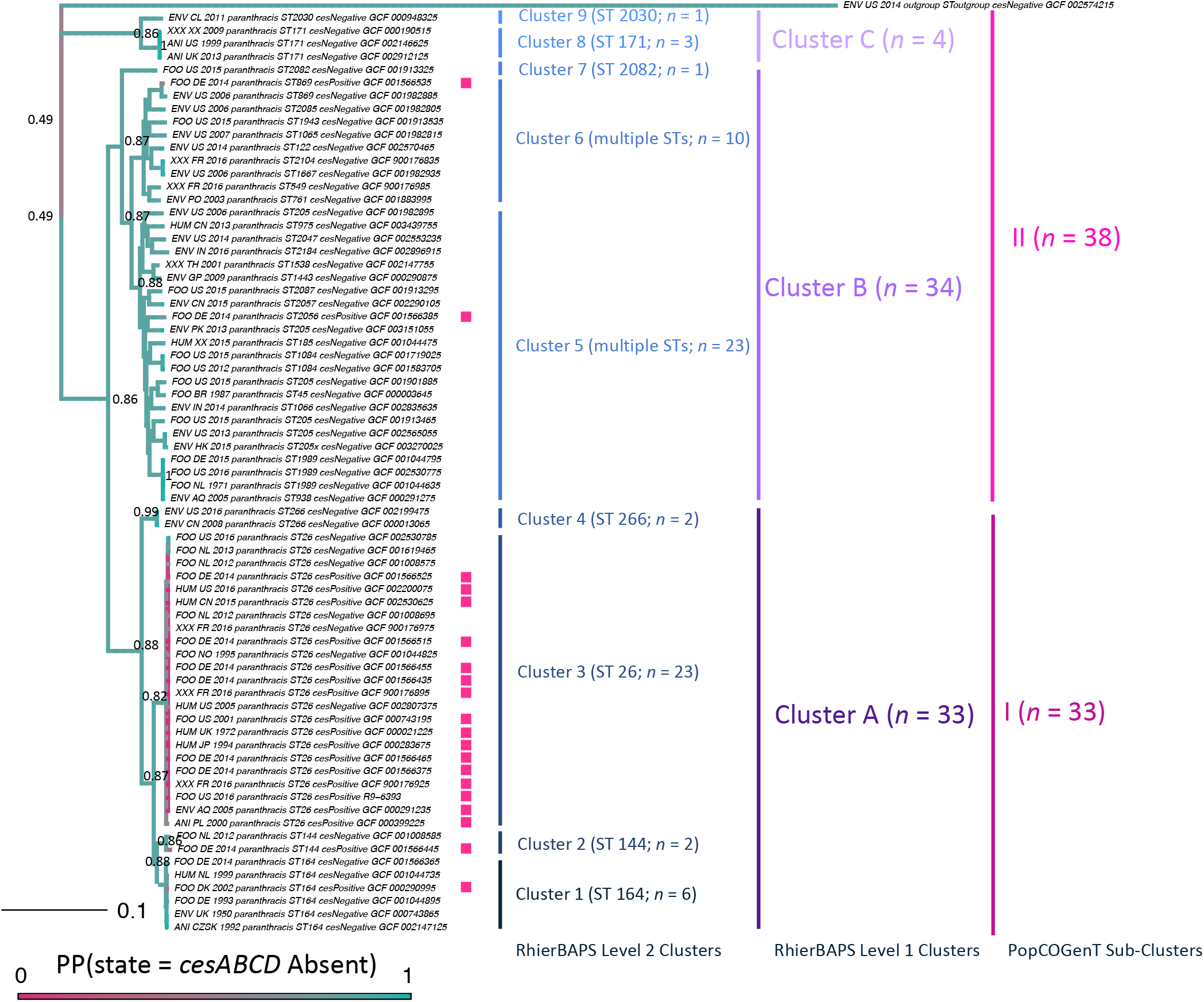
Maximum likelihood phylogeny constructed using core SNPs identified among 71 emetic Group III *B. cereus s.l.* genomes and their closely related, non-emetic counterparts, plus outgroup genome *B. cereus s.l.* str. AFS057383. Tip labels of genomes possessing cereulide synthetase encoding genes *cesABCD* are annotated with a pink square. Clade labels correspond to (i) RhierBAPs level 2 cluster assignments, denoted as Cluster 1 to 9, with number of isolates assigned to a cluster (*n*) and sequence type (ST) determined using *in silico* multi-locus sequence typing (MLST) listed in parentheses; (ii) RhierBAPs level 1 cluster assignments, denoted as Cluster A, B, and C; (iii) PopCOGenT sub-cluster assignments, denoted as I and II. Tree edge and node colors correspond to the posterior probability (PP) of being in a *ces-*negative state, obtained using an empirical Bayes approach, in which a continuous-time reversible Markov model was fitted, followed by 1,000 simulations of stochastic character histories using the fitted model and tree tip states. Equal root node prior probabilities for *ces*-positive and *ces*-negative states were used. Node labels denote selected PP values, chosen for readability. The phylogeny was rooted along the outgroup genome, and branch lengths are reported in substitutions/site.

The 150 Group III genomes queried here (which included all 30 publicly available *ces*-positive genomes, as well as 30 *ces*-positive genomes from a 2016 emetic foodborne outbreak) were distributed into three major clusters and nine sub-clusters using RhierBAPs, with *ces*-positive isolates present in two and five clusters and sub-clusters, respectively (Figure 1). When PopCOGenT was used to delineate populations using recent gene flow, genomes were distributed among two sub-clusters (i.e., populations), with *ces*-positive genomes present in both sub-clusters. All genomes were assigned to a single “main cluster”, a unit that has been proposed to mirror the “species” definition applied to plants and animals (Figure 1) (22). Congruent with these findings, pairwise average nucleotide identity (ANI) values calculated between the 150 genomes confirmed that all cereulide-producing Group III strains would be considered to be members of the same genomospecies using any previously proposed genomospecies threshold for *B. cereus s.l.* (i.e., 92.5-96 ANI) (21, 23–26). However, considerable genomic diversity existed among cereulide-producing isolates, as *ces*-positive genomes could share as low as 97.5 ANI with others (Figure 2).

**Figure 2.**
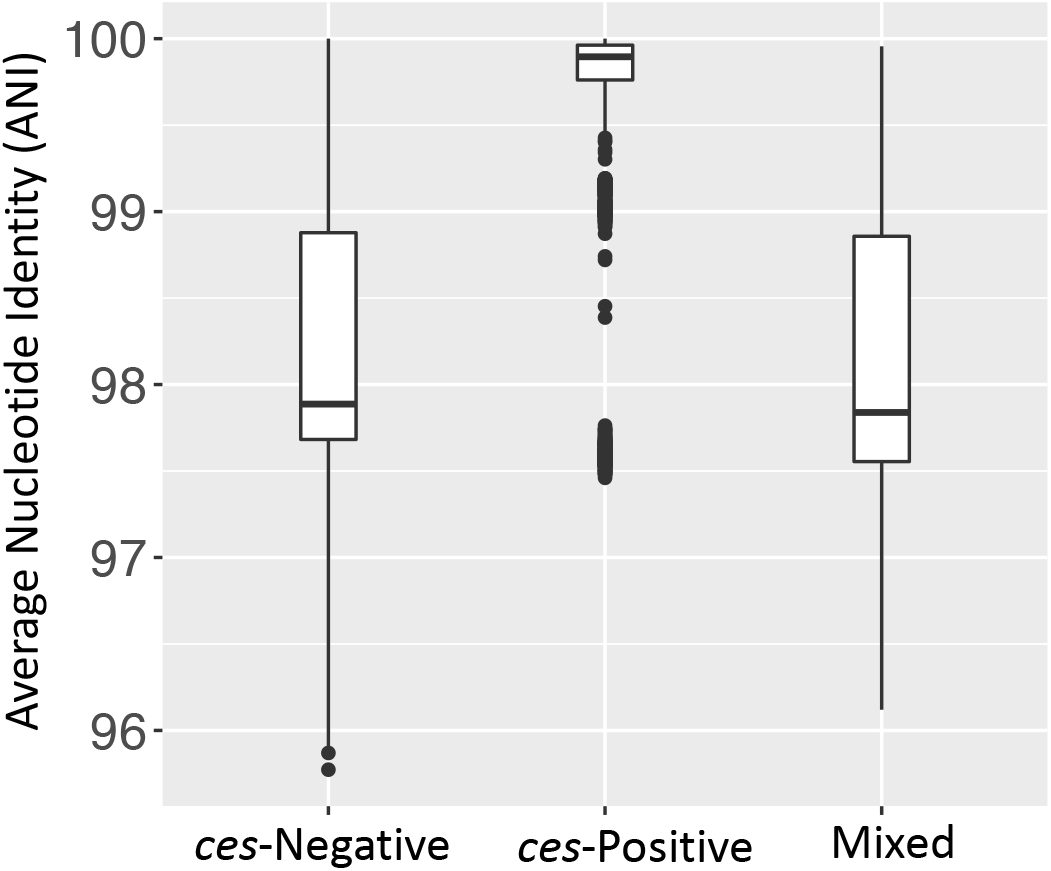
Pairwise average nucleotide identity (ANI) values calculated between Group III *B. cereus s.l.* genomes in which (i) both the query and reference genome lacked *cesABCD* (*ces*-negative; *n =* 90); (ii) both the query and reference genome possessed *cesABCD* (*ces*-positive; *n* = 60); (iii) the query genome possessed *cesABCD* and the reference genome lacked *cesABCD* and vice versa (mixed). Pairwise ANI values were calculated using FastANI version 1.0. Lower and upper box hinges correspond to the first and third quartiles, respectively. Lower and upper whiskers extend from the hinge to the smallest and largest values no more distant than 1.5 times the interquartile range from the hinge, respectively. Points represent pairwise distances that fall beyond the ends of the whiskers.

The common ancestor of all *ces*-positive Group III genomes was predicted to not possess *cesABCD* and, thus, not be capable of cereulide production, regardless of outgroup or use of core or majority SNPs (*ces-*negative state posterior probability [PP] 0.86-0.89; Figure 1, Supplemental Figures S1 and S2, and Supplemental Table S2). For STs 144, 164, 869, and 2056, a single *ces*-positive isolate was present among genomes assigned to the ST (Figure 1). Consequently, a single acquisition event was predicted to be responsible for the presence of *ces*-positive lineages within each of these STs, and the common ancestor shared by each ST encompassing more than one genome was predicted to lack *ces* (Figure 1, Supplemental Figures S1 and S2, and Supplemental Table S2).

### ST 26 first acquired the ability to cause emetic foodborne illness in the twentieth century

ST 26 was the only ST that encompassed multiple *ces*-negative and *ces*-positive strains (Figure 1); therefore, the dynamics of cereulide synthetase loss and gain could be analyzed among members of this lineage. ST 26 isolates in this study were predicted to have evolved from a common ancestor that existed circa 1904 (estimated node age of 1904.3, with a 95% highest posterior density [HPD] interval of 1837.1-1957.8 for common ancestor node heights; Figure 3) with an estimated evolutionary rate of 3.04 × 10^−7^ substitutions/site/year (95% HPD 1.47 × 10^−7^ −4.74 × 10^−7^ substitutions/site/year). Ancestral state reconstruction within ST 26 indicated that the ST 26 common ancestor did not possess cereulide synthetase (*ces*-negative state PP 0.60-0.93; Figure 4, Supplemental Figure S3, and Supplemental Table S2). Rather, *cesABCD* were predicted to have been first acquired within ST 26 between ≈1904 and ≈1931 (95% HPD 1837.1-1957.8 and 1893.2-1959.0 for common ancestor node heights, respectively; Figures 3 and 4 and Supplemental Figure S3). Subsequent losses of *cesABCD* among ST 26 were predicted to have occurred on three occasions: (i) one after 1946 (common ancestor node height 95% HPD 1914.5-1971.0); (ii) one after 1962.9 (common ancestor node height 95% HPD 1938.1-1985.0); and (iii) one between 1961.6 and 1966.7 (95% HPD 1934.7-1983.0 and 1941.6-1987.7, respectively; Figures 3 and 4 and Supplemental Figure S3) (16).

**Figure 3.**
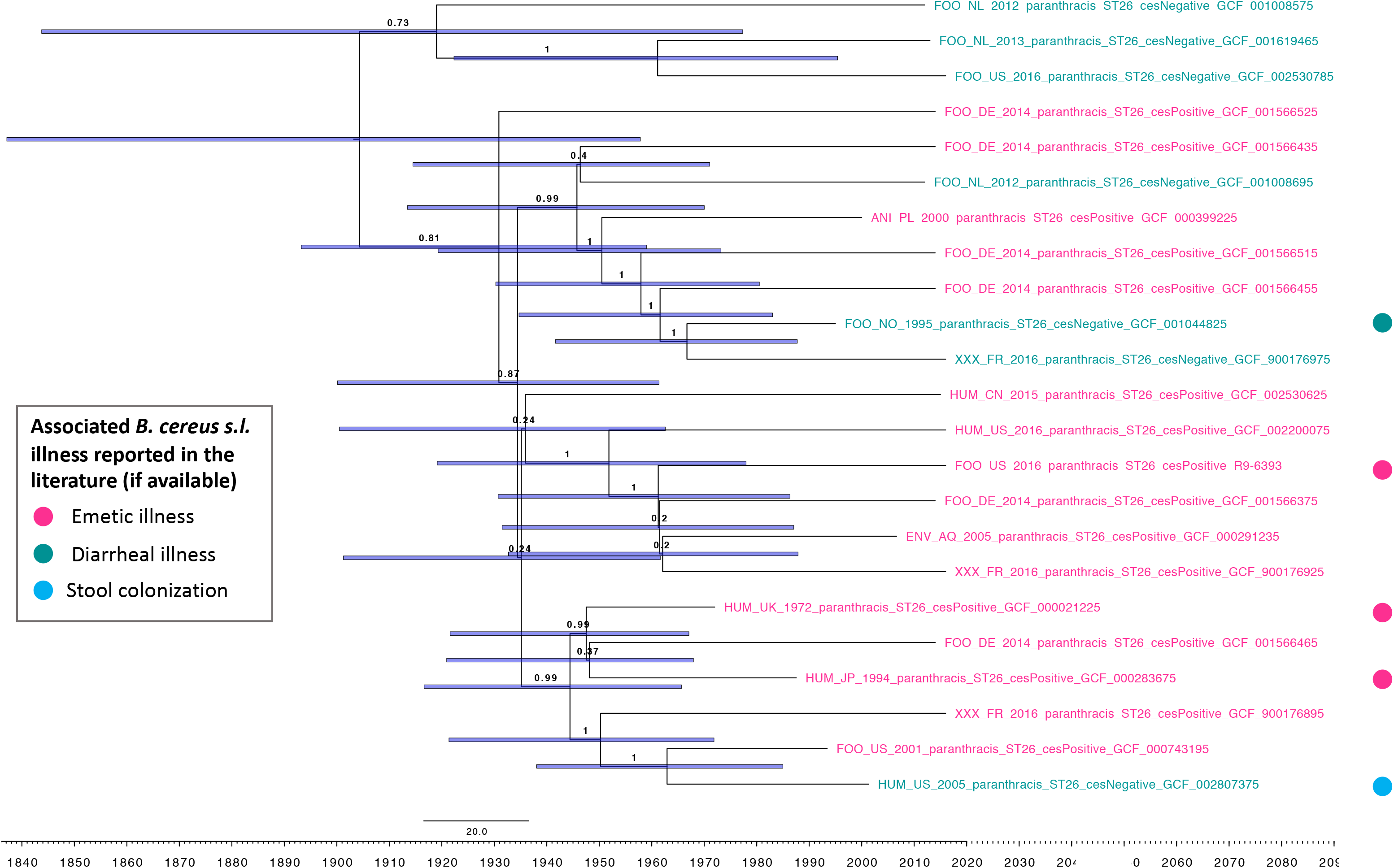
Rooted, time-scaled maximum clade credibility (MCC) phylogeny constructed using core SNPs identified among 23 Group III *B. cereus s.l.* genomes belonging to sequence type (ST) 26. Tip label colors denote *ces*-positive (pink) and *ces*-negative (teal) genomes predicted to be capable and incapable of producing cereulide, respectively. Tip labels of isolates that could be associated with a known *B. cereuss.l.* illness in the literature (emetic, diarrheal, or stool colonization) are annotated on the right side with a pink, teal, or blue circle, respectively (note that additional isolates were associated with illness; however, these are not annotated, as the type of illness could not be confirmed from the available literature). Branch labels denote posterior probabilities of branch support. Time in years is plotted along the X-axis, with branch length reported in substitutions/site/year. Node bars denote 95% highest posterior density (HPD) intervals for common ancestor node heights. Core SNPs were identified using Snippy version 4.3.6. The phylogeny was constructed using the results of five independent runs using a relaxed lognormal clock model, the Standard_TVMef nucleotide substitution model, and the Birth Death Skyline Serial population model implemented in BEAST version 2.5.1, with 10% burn-in applied to each run. LogCombiner-2 was used to combine BEAST 2 log files, and TreeAnnotator-2 was used to construct the phylogeny using common ancestor node heights.

**Figure 4.**
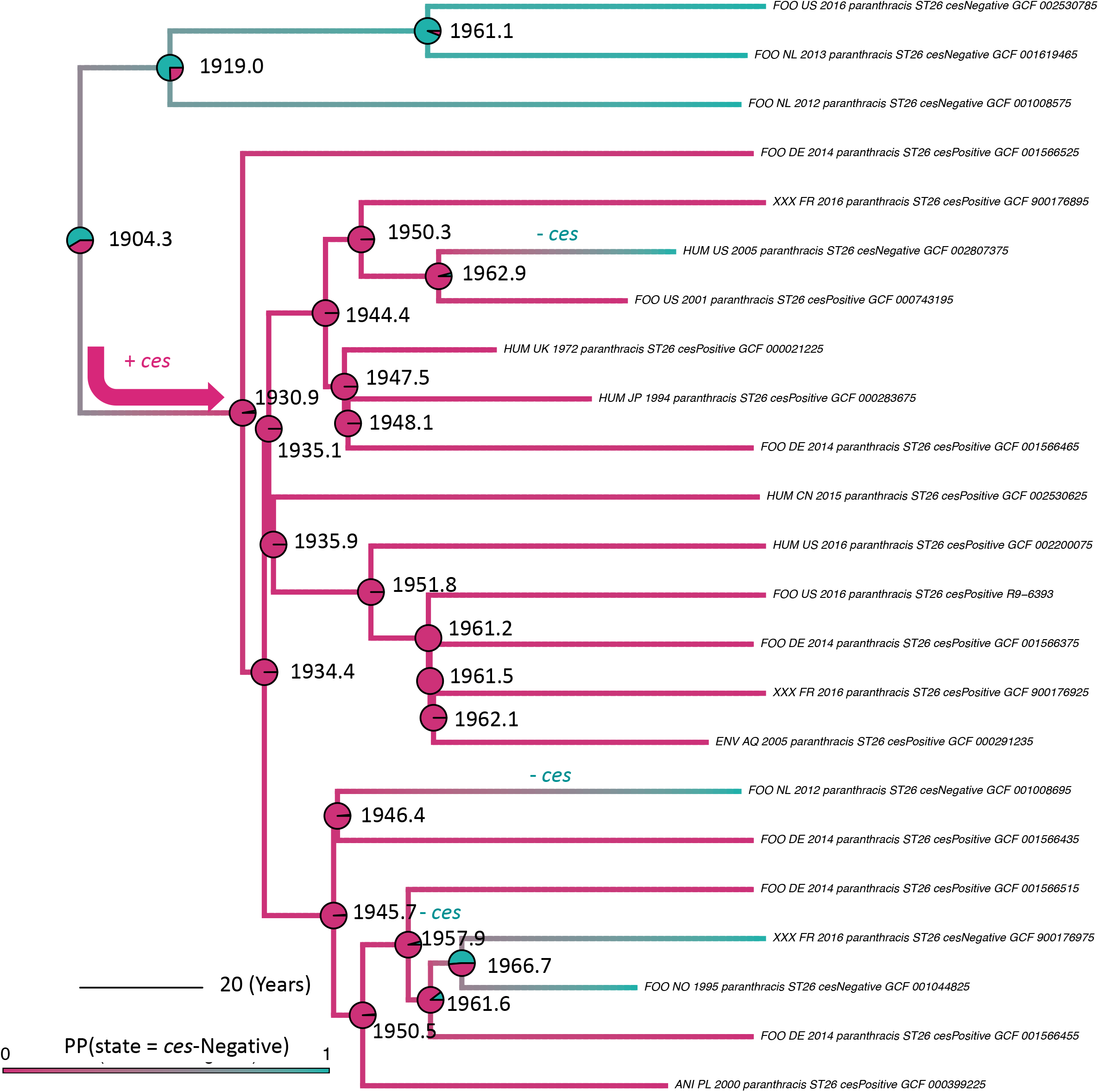
Rooted, time-scaled maximum clade credibility (MCC) phylogeny constructed using core SNPs identified among 23 Group III *B. cereus s.l.* genomes belonging to sequence type (ST) 26. Branch color corresponds to posterior density, denoting the probability of a lineage being in a *ces*-negative state as determined using ancestral state reconstruction. Pie charts at nodes denote the posterior probability (PP) of a node being in a *ces*-negative (teal) or *ces*-positive (pink) state. Arrows along branches denote a *ces* gain event. Labels along branches denote a *ces* gain or loss event (denoted by + *ces* or – *ces*, respectively). Node labels correspond to node ages in years, while branch lengths are reported in substitutions/site/year. Core SNPs were identified using Snippy version 4.3.6. The phylogeny was constructed using the results of five independent runs using a relaxed lognormal clock model, the Standard_TVMef nucleotide substitution model, and the Birth Death Skyline Serial population model implemented in BEAST version 2.5.1, with 10% burn-in applied to each run. LogCombiner-2 was used to combine BEAST 2 log files, and TreeAnnotator-2 was used to construct the phylogeny using common ancestor node heights. Ancestral state reconstruction was performed using a prior on the root node in which the probability of the ST 26 ancestor being *ces*-positive or *ces*-negative was estimated using the make.simmap function in the phytools package in R. For ancestral state reconstruction results obtained using different root node priors, see Supplemental Figure S3.

### Choice of emetic Group III *B. cereus s.l.* reference genome for reference-based SNP calling affects ST 26 phylogenomic topology

SNP identification using reference-based approaches and subsequent phylogeny construction are critical methods used in foodborne pathogen surveillance and outbreak investigation efforts. To determine if choice of emetic reference genome could affect the topology of the ST 26 phylogeny, SNPs were identified among all 64 ST 26 genomes using four reference-based SNP calling pipelines and six emetic reference genomes, which encompassed all observed Group III emetic STs (Table 1). Notably, the emetic Group III genome that was most distantly related to ST 26 (ST 869) did not yield sufficient resolution to produce a phylogeny when it was used as a reference for BactSNP/Gubbins and Snippy/Gubbins (Tables 1 and 2). For the BactSNP pipeline, the emetic ST 2056 genome additionally did not yield an alignment of SNPs among ST 26 isolates when it was used as a reference (Tables 1 and 2).

**Table 1.**
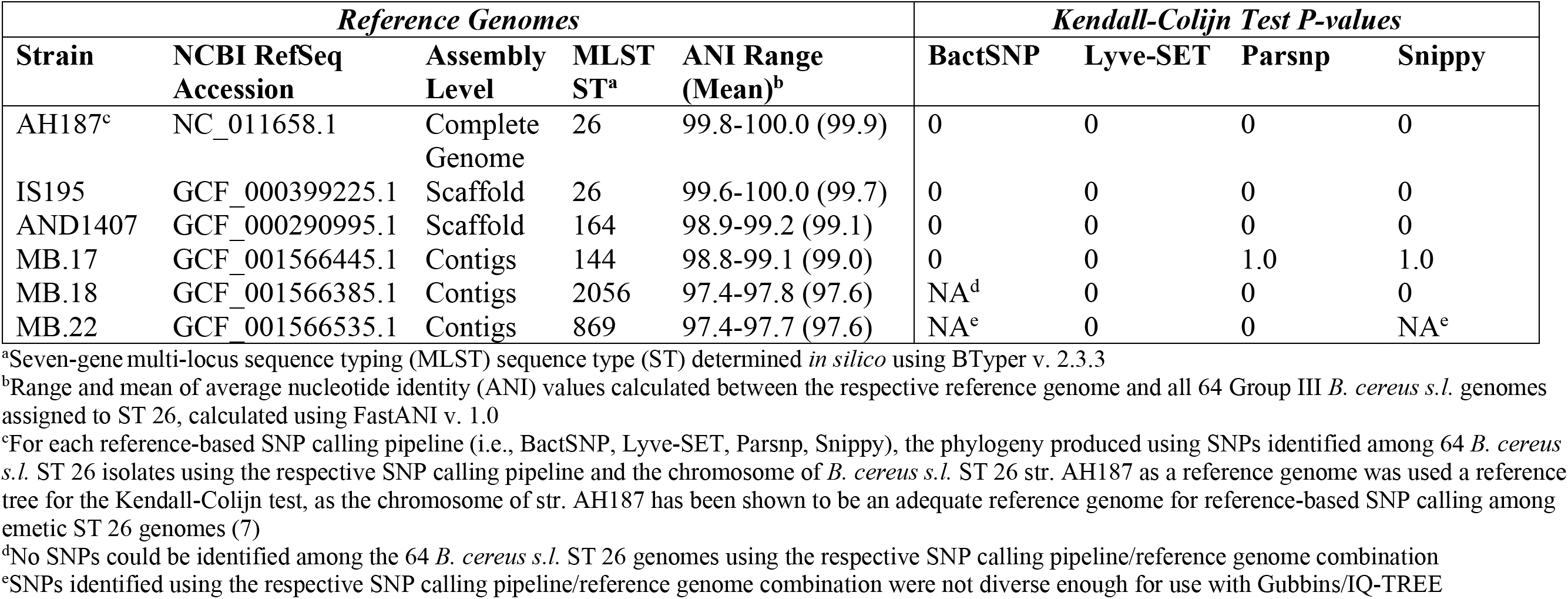
Topological comparisons of *B. cereus s.l.* ST 26 phylogenies constructed using various SNP calling pipeline/reference genome combinations.

**Table 2.**
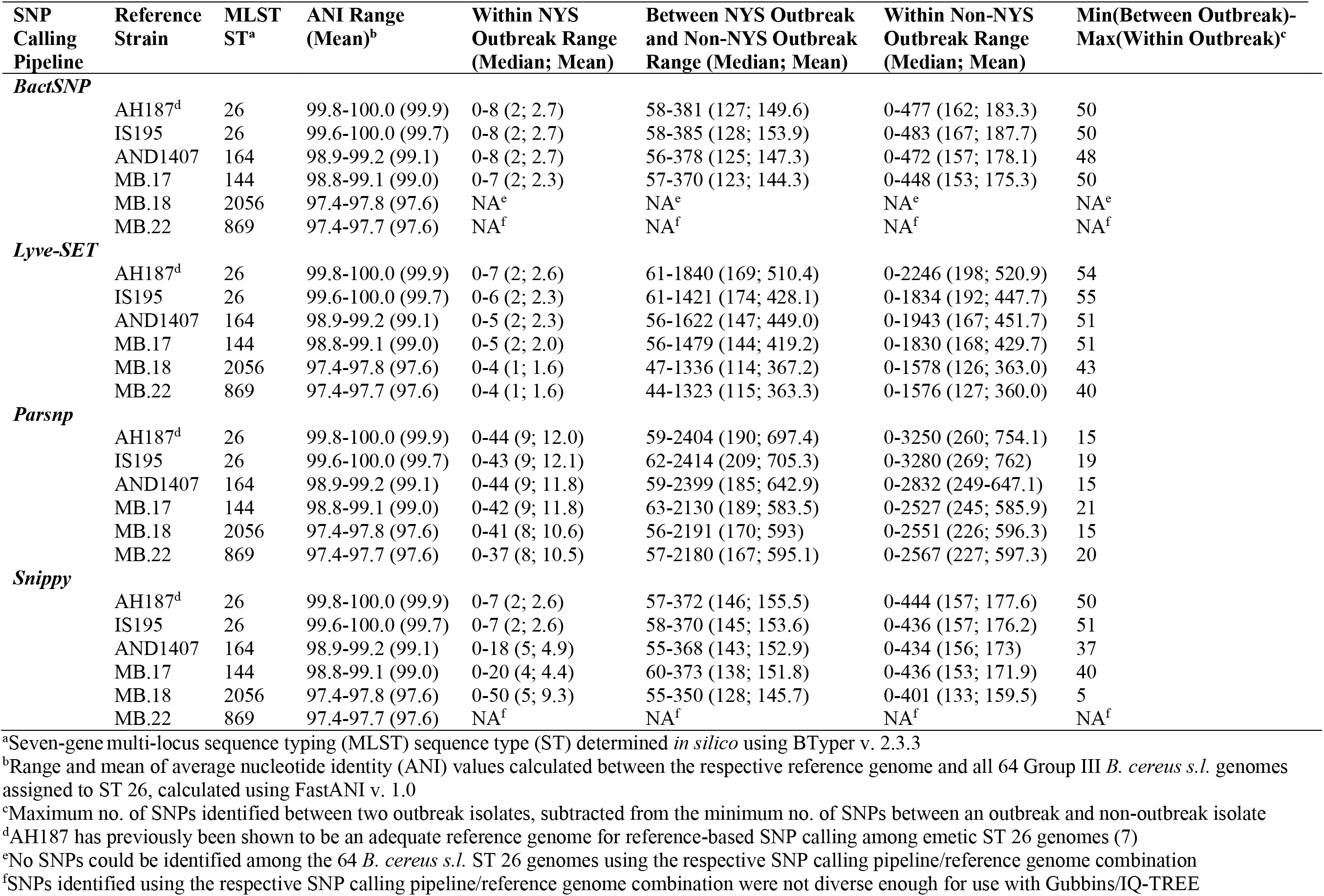
Pairwise SNP differences calculated between 64 *B. cereus s.l.* ST 26 isolates, including 30 e metic isolates from a 2016 foodborne outbreak in New York State (NYS), using various SNP calling pipeline/reference genome combinations.

| SNP Calling Pipeline | Reference Strain | MLST ST<sup>a</sup> | ANI Range (Mean)<sup>b</sup> | Within NYS Outbreak Range (Median; Mean) | Between NYS Outbreak and Non-NYS Outbreak Range (Median; Mean) | Within Non-NYS Outbreak Range (Median; Mean) | Min(Between Outbreak)-Max(Within Outbreak)<sup>c</sup> |
| --- | --- | --- | --- | --- | --- | --- | --- |
| ***BactSNP*** |  |  |  |  |  |  |  |
|  | AH187<sup>d</sup> | 26 | 99.8-100.0 (99.9) | 0-8 (2; 2.7) | 58-381 (127; 149.6) | 0-477 (162; 183.3) | 50 |
|  | IS195 | 26 | 99.6-100.0 (99.7) | 0-8 (2; 2.7) | 58-385 (128; 153.9) | 0-483 (167; 187.7) | 50 |
|  | AND1407 | 164 | 98.9-99.2 (99.1) | 0-8 (2; 2.7) | 56-378 (125; 147.3) | 0-472 (157; 178.1) | 48 |
|  | MB.17 | 144 | 98.8-99.1 (99.0) | 0-7 (2; 2.3) | 57-370 (123; 144.3) | 0-448 (153; 175.3) | 50 |
|  | MB.18 | 2056 | 97.4-97.8 (97.6) | NA<sup>e</sup> | NA<sup>e</sup> | NA<sup>e</sup> | NA<sup>e</sup> |
|  | MB.22 | 869 | 97.4-97.7 (97.6) | NA<sup>f</sup> | NA<sup>f</sup> | NA<sup>f</sup> | NA<sup>f</sup> |
| ***Lyve-SET*** |  |  |  |  |  |  |  |
|  | AH187<sup>d</sup> | 26 | 99.8-100.0 (99.9) | 0-7 (2; 2.6) | 61-1840 (169; 510.4) | 0-2246 (198; 520.9) | 54 |
|  | IS195 | 26 | 99.6-100.0 (99.7) | 0-6 (2; 2.3) | 61-1421 (174; 428.1) | 0-1834 (192; 447.7) | 55 |
|  | AND1407 | 164 | 98.9-99.2 (99.1) | 0-5 (2; 2.3) | 56-1622 (147; 449.0) | 0-1943 (167; 451.7) | 51 |
|  | MB.17 | 144 | 98.8-99.1 (99.0) | 0-5 (2; 2.0) | 56-1479 (144; 419.2) | 0-1830 (168; 429.7) | 51 |
|  | MB.18 | 2056 | 97.4-97.8 (97.6) | 0-4 (1; 1.6) | 47-1336 (114; 367.2) | 0-1578 (126; 363.0) | 43 |
|  | MB.22 | 869 | 97.4-97.7 (97.6) | 0-4 (1; 1.6) | 44-1323 (115; 363.3) | 0-1576 (127; 360.0) | 40 |
| ***Parsnp*** |  |  |  |  |  |  |  |
|  | AH187<sup>d</sup> | 26 | 99.8-100.0 (99.9) | 0-44 (9; 12.0) | 59-2404 (190; 697.4) | 0-3250 (260; 754.1) | 15 |
|  | IS195 | 26 | 99.6-100.0 (99.7) | 0-43 (9; 12.1) | 62-2414 (209; 705.3) | 0-3280 (269; 762) | 19 |
|  | AND1407 | 164 | 98.9-99.2 (99.1) | 0-44 (9; 11.8) | 59-2399 (185; 642.9) | 0-2832 (249-647.1) | 15 |
|  | MB.17 | 144 | 98.8-99.1 (99.0) | 0-42 (9; 11.8) | 63-2130 (189; 583.5) | 0-2527 (245; 585.9) | 21 |
|  | MB.18 | 2056 | 97.4-97.8 (97.6) | 0-41 (8; 10.6) | 56-2191 (170; 593) | 0-2551 (226; 596.3) | 15 |
|  | MB.22 | 869 | 97.4-97.7 (97.6) | 0-37 (8; 10.5) | 57-2180 (167; 595.1) | 0-2567 (227; 597.3) | 20 |
| ***Snippy*** |  |  |  |  |  |  |  |
|  | AH187<sup>d</sup> | 26 | 99.8-100.0 (99.9) | 0-7 (2; 2.6) | 57-372 (146; 155.5) | 0-444 (157; 177.6) | 50 |
|  | IS195 | 26 | 99.6-100.0 (99.7) | 0-7 (2; 2.6) | 58-370 (145; 153.6) | 0-436 (157; 176.2) | 51 |
|  | AND1407 | 164 | 98.9-99.2 (99.1) | 0-18 (5; 4.9) | 55-368 (143; 152.9) | 0-434 (156; 173) | 37 |
|  | MB.17 | 144 | 98.8-99.1 (99.0) | 0-20 (4; 4.4) | 60-373 (138; 151.8) | 0-436 (153; 171.9) | 40 |
|  | MB.18 | 2056 | 97.4-97.8 (97.6) | 0-50 (5; 9.3) | 55-350 (128; 145.7) | 0-401 (133; 159.5) | 5 |
|  | MB.22 | 869 | 97.4-97.7 (97.6) | NA<sup>f</sup> | NA<sup>f</sup> | NA<sup>f</sup> | NA<sup>f</sup> |

<sup>a</sup>Seven-gene multi-locus sequence typing (MLST) sequence type (ST) determined *in silico* using BTypyer v. 2.3.3
<sup>b</sup>Range and mean of average nucleotide identity (ANI) values calculated between the respective reference genome and all 64 Group III *B. cereus s.l.* genomes assigned to ST 26, calculated using FastANI v. 1.0
<sup>c</sup>Maximum no. of SNPs identified between two outbreak isolates, subtracted from the minimum no. of SNPs between an outbreak and non-outbreak isolate
<sup>d</sup>AH187 has previously been shown to be an adequate reference genome for reference-based SNP calling among emetic ST 26 genomes (7)
<sup>e</sup>No SNPs could be identified among the 64 *B. cereus s.l.* ST 26 genomes using the respective SNP calling pipeline/reference genome combination
<sup>f</sup>SNPs identified using the respective SNP calling pipeline/reference genome combination were not diverse enough for use with Gubbins/IQ-TREE

For the remaining SNP calling pipeline/reference genome combinations, the resulting phylogeny was compared to the phylogeny produced using the respective pipeline and the chromosome of ST 26 str. AH187 as a reference. In addition to being a well-characterized emetic strain for which a closed genome is available, str. AH187 was closely related to the 64 ST 26 isolates queried here and has previously been shown to serve as an adequate reference genome for SNP calling within ST 26 (7). For all SNP calling pipelines, phylogenies produced using the genomes of emetic ST 26 str. IS195 and emetic ST 164 str. AND1407 as references were more topologically similar to those produced using str. AH187 than would be expected by chance (Kendall-Colijn *P* < 0.05 after a Bonferroni correction; Table 1). However, the topology of phylogenies produced using Parsnp and Snippy with emetic ST 144 str. MB.17 differed from that produced using str. AH187 (Kendall-Colijn *P* > 0.05 after a Bonferroni correction; Table 1). Lyve-SET was the only pipeline that produced phylogenies that were more topologically similar to that produced using str. AH187 than would be expected by chance, regardless of emetic reference (Kendall-Colijn *P* < 0.05 after a Bonferroni correction; Table 1).

Despite producing phylogenies that resembled the AH187 phylogeny for five of six emetic reference genomes (Kendall-Colijn *P* < 0.05 after a Bonferroni correction; Table 1), core SNP alignments produced with Parsnp yielded relatively large pairwise SNP distances between emetic ST 26 genomes from a known outbreak (7). Regardless of reference genome selection, the difference between the minimum number of SNPs shared between outbreak and non-outbreak isolates and the maximum number of SNPs detected between two outbreak isolates was less than the maximum number of SNPs shared between two outbreak isolates (Table 2). A similar phenomenon was observed when Snippy was used with a distant emetic ST 2056 strain as a reference (Table 2).

## Discussion

### Group III *B. cereus s.l.* isolates capable of causing emetic foodborne illness are not clonal

Cereulide-producing *B. cereus s.l.* strains are responsible or suspected to be responsible for thousands of cases of foodborne illness each year worldwide (2), including rare but severe forms of illness which may result in death (27–31). While efforts to characterize this important pathogen using whole-genome sequencing have begun only recently, the amount of publicly available genomic data derived from “emetic *B. cereus*” has been increasing (21). Consequently, the current dogma regarding the evolutionary history of this group of organisms must be revisited; while prior studies assert that cereulide-producing Group III members represent a highly clonal complex within *B. cereus s.l.* (10, 16), other efforts have hinted that “emetic *B. cereus*” showcases a considerable degree of genomic diversity (21, 32–34).

Using all publicly available emetic Group III *B. cereus s.l.* genomes and the non-emetic genomes interspersed among them, we show on a whole-genome scale that “emetic *B. cereus*” is not clonal. Emetic toxin production capabilities within Group III are not the result of a single cereulide synthetase gain event followed by subsequent proliferation; rather, the common ancestor of all cereulide-producing Group III isolates was likely incapable of producing cereulide, and emetic toxin production capabilities resulted from at least five independent cereulide synthetase acquisition events (at least one in each of STs 26, 144, 164, 869, and 2056; Figures 1 and 4). Pairwise ANI values calculated between emetic Group III strains were as low as 97.5 ANI; for comparison, all members of the highly similar *B. anthracis* lineage commonly attributed to anthrax illness share ≥99.9 ANI with one another (21, 35), while genomes belonging to *Salmonella enterica* subspecies *enterica* (which is not considered to be clonal) can share pairwise ANI values as low as 97.0 (calculated between 425 genomes described by Worley, et al., using FastANI v. 1.0 as described in the Methods section) (36).

These findings are important, as unexpected diversity can confound bioinformatic analyses used to identify outbreaks from genomic data. For example, an evolutionarily distant reference genome can affect which SNPs are identified during reference-based SNP calling among bacterial genomes (7, 37–40). This can, in turn, affect metrics used to determine whether an isolate should be included or excluded from an outbreak (e.g., the topology of a resulting phylogeny, pairwise SNP cut-offs) (7, 38–40). Here, we showed that emetic Group III isolates are considerably diverse, so much so that the use of some “emetic *B. cereus*” genomes as references for SNP calling can lead to a topologically confounding loss of resolution. The use of BactSNP/Gubbins and Snippy/Gubbins with distant emetic ST 869 as a reference, for example, yielded SNPs that could not reliably differentiate ST 26 genomes from each other. In an outbreak scenario, these approaches would incorrectly place non-outbreak isolates among outbreak ones, potentially confounding an investigation. It is thus essential that the diversity of “emetic *B. cereus*” is acknowledged and accounted for to ensure that epidemiological investigations are not hindered.

### One pathogen, two illnesses: ST 26 *B. cereus s.l.* has oscillated between “emetic” and “diarrheal” foodborne pathogen throughout the twentieth century

“*B. cereus*” was first established as the causative agent of a diarrheal form of foodborne illness in the 1950s (20, 41). Notably, prior to the 1970s, illnesses attributed to “*B. cereus*” were of the diarrheal type (i.e., toxicoinfection characterized by symptoms of watery diarrhea that onset 8-16 h after ingestion) (20). However, in the 1970s, a novel type of “*B. cereus*” illness, emetic intoxication, began to be reported (20). Characterized by symptoms of vomiting and nausea and a relatively short incubation time (i.e., 0.5-6 h), “*B. cereus*” emetic illness was first described in the United Kingdom in 1971, and was linked to the consumption of rice served at restaurants and take-away outlets (20). It has been hypothesized that emetic toxin production may confer a selective advantage (16), and the results reported here support the hypothesis that cereulide synthetase was acquired by some Group III lineages relatively recently in their evolutionary histories (16). Here, we show that ST 26, which has frequently been associated with emetic foodborne illness (7, 32, 42, 43), first acquired cereulide synthetase and, thus, the ability to cause emetic illness in the twentieth century, likely between 1904 and 1931 (95% HPD interval of 1837.1-1959.0). This indicates that cereulide-producing *B. cereus s.l.* may have been responsible for cryptic cases of emetic intoxication prior to the 1970s; however, it is unsurprising that these cases would go undetected or unattributed to *B. cereus s.l.*, due to the mild and transient symptoms typically associated with this illness (1, 44).

The temporal characterization of cereulide synthetase acquisition and loss provided here additionally showcases that ST 26 has transitioned between an emetic and non-emetic pathogen over the course of its evolutionary history. This is important, as *ces*-negative members of ST 26 still present a relevant public health and food safety risk, as they may still be capable of causing diarrheal illness. For example, the lineage to which ST 26 str. NVH 0075-95 belongs lost *ces* between ≈1962 and 1967. While previously shown to be incapable of producing cereulide, this strain produces diarrheal non-hemolytic enterotoxin (Nhe), is highly cytotoxic, and was isolated from vegetable stew associated with a diarrheal outbreak in Norway (16, 45, 46). Additionally, cereulide-producing strains can be high producers of diarrheal enterotoxins (8). It has been hypothesized that the simultaneous ingestion of food contaminated with cereulide alongside the cereulide- and enterotoxin-producing strains themselves may be responsible for a mixture of diarrheal and emetic symptoms among some cases of *B. cereus s.l.* foodborne illness (8), and this may partially explain why these illnesses may not always present within a strictly “emetic-vs-diarrheal” dichotomy (7, 8).

### Heterogeneous emetic phenotype presentation among diverse Group III *B. cereus s.l.* isolates can yield taxonomic inconsistencies: the “emetic *B. cereus*” problem

Recent inconsistencies have arisen in the *B. cereus s.l.* taxonomic space: *B. paranthracis*, a novel species proposed in 2017 (26), was found to encompass all cereulide-producing Group III *B. cereus s.l.* strains at conventional species thresholds (21). Using multiple metrics for species delineation (i.e., ANI-based genomospecies assignment, methods querying recent gene flow), we confirm that all cereulide-producing Group III isolates, along with *B. paranthracis* and the other *ces*-negative isolates queried here (excluding outgroup genomes), belong to a single genomospecies. However, using “*B. paranthracis*” to describe cereulide-producing Group III members is problematic, as *B. paranthracis* was only recently proposed as a novel species, is not well-recognized outside the small *B. cereus s.l.* taxonomic space, and hence would not typically be equated with a foodborne pathogen (21).

Referring to cereulide-producing Group III lineages as “emetic *B. cereus*”, however, is also problematic. Because cereulide synthetase is often plasmid-encoded (1, 9, 10, 47), it may be possible for emetic toxin production capabilities to be lost, gained, present across multiple lineages, and absent within individual lineages (21). Here we show that this is not just a hypothetical scenario: even with the limited number of genomes presently available, we observed five cereulide synthetase gain events across Group III, and three loss events within ST 26 alone, indicating that cereulide synthetase loss and gain is a dynamic and ongoing process. Additionally, a taxonomic label of “*B. cereus*” as it is applied to Group III *B. cereus s.l.* is misleading, as Group III strains are not actually members of the *B. cereus sensu stricto* (*s.s.*) species, regardless of which previously proposed genomospecies threshold for *B. cereus s.l.* is used to define species (i.e., 92.5-96 ANI) (7, 21, 23–26).

Taxonomic labels used to refer to *ces*-negative isolates interspersed among cereulide-producing Group III isolates (i.e., the *ces*-negative isolates queried here) are even more ambiguous. Some of these *ces*-negative isolates are capable of causing diarrheal illness (16, 45, 46) and are thus relevant threats to global public health; however, prior to 2020, there was no standardized nomenclature with which these isolates could be described. For example, the following names have been used to refer to *ces*-negative, Group III strains: (i) “emetic-like *B. cereus*”, (ii) “*B. cereus*”, (iii) “Group III *B. cereus*”, (iv) “*B. paranthracis*”, or (v) “*B. cereus sensu stricto*”/“*B. cereus s.s*”, although it should be noted that *B. cereus s.s.* is a misnomer; as mentioned previously, Group III strains do not fall within the genomospecies boundary of the *B.cereus s.s.* type strain and thus are not actually members of the *B. cereus s.s.* species (12, 16, 26, 48–51).

It is thus essential that microbiologists, clinicians, public health officials, and industrial professionals find common ground and adhere to a standardized nomenclature when describing Group III *B. cereus s.l.* Recently, we have proposed a taxonomic framework which can account for emetic heterogeneity among *B. cereus s.l.* genomes through the incorporation of a standardized collection of biovar terms (21), including the biovar term “Emeticus”. Using this framework, all cereulide-producing members of *B. cereus s.l.* (including “emetic *B. weihenstephanensis*”) can be referenced using the name *B.* Emeticus. All cereulide-producing Group III lineages are *B. mosaicus* subspecies *cereus* biovar Emeticus (full name) or *B. cereus* biovar Emeticus (shorted subspecies notation), while the *ces*-negative isolates interspersed among them are *B. mosaicus* subsp. *cereus* (full name) or *B. cereus* (shortened subspecies notation) (21). Note that “*sensu stricto*” or “*s.s.*” is not appended to these names; as mentioned above, Group III *B. cereus s.l.* lineages do not belong to the same species as Group IV *B. cereus s.s.* type strain ATCC 14579 (7, 21).

This study is the first to offer insight into the temporal dynamics of cereulide synthetase loss and gain among Group III *B. cereus s.l.*, and it showcases the importance of accounting for emetic heterogeneity among Group III lineages. As genomic sequencing grows in popularity and more Group III genomes are sequenced, the estimates provided here can be further refined and improved. Furthermore, it is likely that additional cereulide synthetase loss and gain events will be observed, and that previously uncharacterized emetic Group III lineages will be discovered.

## Methods

### Acquisition of Group III *B. cereus s.l.* genomes and metadata

All genomes submitted to NCBI RefSeq (52) as a published *B. cereus s.l.* species (21, 23–26, 53) were downloaded (*n* = 2,231; accessed November 19, 2018). The ANI function in BTyper v. 2.3.3 (13) was used to calculate ANI values between each genome and the type strain/species reference genomes of each of the 18 published *B. cereus s.l.* species as they existed in 2019 (7). Genomes that (i) most closely resembled *B. paranthracis* and (ii) shared an ANI value ≥ 95 with *B. paranthracis* were used in subsequent steps (*n* = 120), as this set of genomes contained all Group III genomes that possessed genes encoding cereulide synthetase (described in detail below). These genomes were supplemented with 30 genomes of strains isolated in conjunction with a 2016 emetic outbreak (7), resulting in 150 Group III *B. cereus s.l.* genomes (Supplemental Table S1). FastANI v. 1.0 (35) was used to confirm that all 150 genomes (i) shared ≥ 95 ANI with the *B. paranthracis* type strain genome, and (ii) most closely resembled the *B. paranthracis* type strain genome when compared to the 18 *B. cereus s.l.* type strain/reference genomes.

Metadata for each of the 150 genomes were obtained using publicly available records, and BTyper was used to assign each genome to a ST using the seven-gene MLST scheme available in PubMLST (Supplemental Text) (54). To assess the emetic potential of each genome, BTyper was used to detect cereulide synthetase genes *cesABCD* in each genome, first using the default coverage and identity thresholds (70 and 50%, respectively), and a second time with 0% coverage to confirm that *cesABCD* were absent from genomes in which they were not detected (the only genome affected by this was one of the outbreak isolates, FSL R9-6384, which had *cesD* split on two contigs). Isolates in which *cesABCD* were not detected were given a designation of *ces*-negative. BTyper was additionally used to detect *cesABCD* in each of the 2,111 *B. cereus s.l.* genomes not included in this study, as well as to assign all genomes to a *panC* group using the typing scheme described by Guinebretiere, et al (12). All 150 genomes selected for this study were assigned to *panC* Group III, and all Group III genomes possessing *cesABCD* were confirmed to have been included in this study. The only other genomes that possessed *cesABCD* belonged to *panC* Group VI and most closely resembled *B. mycoides*/*B. weihenstephanensis* (i.e., “emetic *B. weihenstephanensis*”) (21).

### Construction of Group III *B. cereus s.l.* maximum likelihood phylogenies and ancestral state reconstruction

kSNP3 v. 3.1 (55, 56) was used to identify (i) core and (ii) majority SNPs among the 150 genomes described above, plus one of two outgroup genomes (to ensure that choice of outgroup did not affect ancestral state reconstruction; Supplemental Text), using the optimal *k*-mer size determined by Kchooser (*k* = 21 for both). For each of the four SNP alignments (i.e., each combination of outgroup and either core or majority SNPs), IQ-TREE v.1.6.10 (57–60) was used to construct a maximum likelihood (ML) phylogeny (Supplemental Text).

To ensure that ancestral state reconstruction would not be affected by genomes over-represented in RefSeq (e.g., genomes confirmed or predicted to have been derived from strains isolated from the same outbreak), potential duplicate genomes were removed using isolate metadata and by assessing clustering in the phylogenies described above. One representative genome was selected from clusters that likely consisted of duplicate genomes and/or isolates derived from the same source. For example, this procedure reduced 30 closely related isolates from an outbreak (7) to one isolate. Overall, this approach yielded a reduced, de-replicated set of 71 genomes (Supplemental Table S1). kSNP3 and IQ-TREE were again used to identify core and majority SNPs and construct ML phylogenies among the set of 71 de-replicated genomes, plus each of the two outgroup genomes, as described above, but with *k* adjusted to the optimal *k*-mer size produced by Kchooser (*k* = 23 for both).

To estimate ancestral character states of internal nodes in the Group III phylogeny as they related to cereulide production (i.e., whether a node represented an ancestor that was *ces*-positive or *ces*-negative), the presence or absence of *ces* within each genome was treated as a binary state. Each of the four phylogenies constructed using the de-replicated set of 71 genomes as described above was rooted at its respective outgroup, and stochastic character maps were simulated on each phylogeny using the make.simmap function in the phytools package (61), the all-rates-different (ARD) model, and one of two root node priors (eight total combinations of two root node priors and four phylogenies; Supplemental Text and Supplemental Table S2).

### Assessment of Group III *B. cereus s.l.* population structure

Core SNPs detected among the 71 de-replicated Group III genomes using kSNP3 (see section “Construction of Group III *B. cereus s.l.* maximum likelihood phylogenies and ancestral state reconstruction” above) were used as input for RhierBAPS (62) to identify clusters, using two levels. The same set of 71 genomes was used as input for PopCOGenT (downloaded October 5, 2019) to identify gene flow units and populations (Supplemental Text) (22).

### Construction of Group III *B. cereus s.l.* ST 26 temporal phylogeny

Snippy v. 4.3.6 (63) was used to identify core SNPs among the de-replicated set of 23 ST 26 genomes (see section “Construction of Group III *B. cereus s.l.* maximum likelihood phylogenies and ancestral state reconstruction” above), using the closed chromosome of emetic ST 26 str. AH187 (NCBI RefSeq Assession NC_011658.1) as a reference genome (Supplemental Text). Gubbins v. 2.3.4 (64) was used to remove recombination from the resulting alignment, and snp-sites (65) was used to obtain core SNPs among the 23 genomes. IQ-TREE was used to construct a phylogeny (Supplemental Text), and the temporal signal of the resulting ML phylogeny was assessed using TempEst v. 1.5.3 (*R*^2^ = 0.26 using the best-fitting root) (66).

Using the ST 26 core SNP alignment as input, BEAST v. 2.5.1 (67, 68) was used to construct a tip-dated phylogeny (Supplemental Text). The Standard_TVMef nucleotide substitution model implemented in the SSM package (69) was used with 5 Gamma categories, and an ascertainment bias correction was applied to account for the use of solely variant sites (Supplemental Text) (70). A relaxed lognormal molecular clock (71) was used with an initial clock rate of 1.0 × 10^−9^ substitutions/site/year, and a broad lognormal prior was placed on the ucldMean parameter (in real space, M = 1.0 × 10^−3^ and S = 4.0) (Supplemental Text). A serial Birth-Death Skyline population model (72) was used to account for potential sampling biases stemming from the overrepresentation of strains isolated in recent years (Supplemental Text).

Five independent runs using the model described above were performed, using chain lengths of at least 100 million generations, sampling every 10,000 generations. For each independent replicate, Tracer v. 1.7.1 (73) was used to ensure that each parameter had mixed adequately with 10% burn-in, and LogCombiner-2 was used to combine log and tree files from each independent run (Supplemental Text). TreeAnnotator-2 (74) was used to produce a maximum clade credibility tree from the combined tree files, using Common Ancestor node heights (Supplemental Text).

### Cereulide synthetase ancestral state reconstruction for ST 26 genomes

Ancestral state reconstruction as it related to cereulide production was performed using the temporal ST 26 phylogeny as input (see section “Construction of Group III *B. cereus s.l.* ST 26 temporal phylogeny” above). Stochastic character maps were simulated on the phylogeny using the make.simmap function, the ARD model, and one of three priors on the root node (Supplemental Text).

### Evaluation of the influence of reference genome selection on ST 26 phylogenomic topology

To determine if choice of reference genome affected ST 26 phylogenomic topology, SNPs were identified among all 64 ST 26 genomes using four different reference-based SNP calling pipelines, chosen for their ability to utilize assembled genomes or both assembled genomes and Illumina reads as input: (i) BactSNP v. 1.1.0 (75), (ii) Lyve-SET v. 1.1.4g (76), (iii) Parsnp v. 1.2 (77), and (iv) Snippy v. 4.3.6. For alignments produced using BactSNP and Snippy, Gubbins v.2.3.4 (64) was used to filter out recombination events; for Parsnp, PhiPack (78) was used to remove recombination (Supplemental Text).

Each of four SNP calling pipelines was run six separate times, each time using one of six emetic Group III reference genomes (Table 1 and Supplemental Text). The tested reference genomes represented all available Group III STs in which *cesABCD* were detected. For each SNP calling pipeline, the phylogeny constructed using SNPs identified with emetic ST 26 str. AH187 as a reference genome was treated as a reference tree, as this genome was closely related to all ST 26 isolates in the study and has previously been shown to serve as an adequate reference genome for ST 26 (7). For each of the four SNP calling pipelines, the Kendall-Colijn (79, 80) test described by Katz et al. (76) was used to compare the topology of each tree to the pipeline’s respective AH187 reference phylogeny, using midpoint-rooted trees, a lambda value of 0 (to give weight to tree topology, rather than branch lengths), and a background distribution of 100,000 random trees (Supplemental Text) (76). Pairs of trees were considered to be more topologically similar than would be expected by chance (76) if a significant *P*-value resulted after a Bonferroni correction was applied (*P* < 0.05).

## Data availability

Accession numbers for all isolates included in this study are available in Supplemental Table S1. The raw BEAST 2 XML file, the code used to perform ancestral state reconstruction, and all phylogenies are available at: https://github.com/lmc297/Group_III_bacillus_cereus.

## Acknowledgments

This material is based on work supported by the National Science Foundation Graduate Research Fellowship Program under grant no. DGE-1650441. The work was also partially supported by USDA NIFA grant 2019-67017-29591. The authors would like to acknowledge those who have generously collected and provided the publicly available genomic data and/or metadata used in this study (14, 18, 26, 45, 81–118).

## SUPPLEMENTAL MATERIAL LEGENDS

**Supplemental Figure S1.** Maximum likelihood phylogenies of 71 emetic Group III *B. cereus s.l.* genomes and their closely related, non- emetic counterparts, plus outgroup genomes (A and B) *B. anthracis* str. Ames, and (C) *B. cereus s.l.* str. AFS057383. Phylogenies were constructed using (A) core, and (B and C) majority SNPs. Tree edge and node colors correspond to the posterior probability (PP) of being in a *ces-*negative state, obtained using an empirical Bayes approach, in which a continuous-time reversible Markov model was fitted, followed by 1,000 simulations of stochastic character histories using the fitted model and tree tip states. Equal root node prior probabilities for *ces*-positive and *ces*-negative states were used. Each phylogeny was rooted along its respective outgroup genome, and branch lengths are reported in substitutions/site.

**Supplemental Figure S2.** Maximum likelihood phylogenies of 71 emetic Group III *B. cereus s.l.* genomes and their closely related, non-emetic counterparts, plus outgroup genomes (A) *B. anthracis* str. Ames, and (B) *B. cereus s.l.* str. AFS057383. Phylogenies were constructed using(1) core, and (2) majority SNPs. Tree edge and node colors correspond to the posterior probability (PP) of being in a *ces-*negative state, obtained using an empirical Bayes approach, in which a continuous-time reversible Markov model was fitted, followed by 1,000 simulations of stochastic character histories using the fitted model and tree tip states. Root node prior probabilities for *ces*-positive and *ces*-negative states were estimated using the make.simmap function in the phytools package in R. Each phylogeny is rooted along its respective outgroup, and branch lengths are reported in substitutions/site.

**Supplemental Figure S3.** Rooted, time-scaled maximum clade credibility (MCC) phylogenies constructed using core SNPs identified among 23 Group III *B. cereus s.l.* genomes belonging to sequence type (ST) 26. Ancestral state reconstruction was performed using the following priors on the root node: (A) probability of the root node belonging to a *ces*-positive or *ces*-negative state set to 0.5 each; or (B) probability of the root node being in a *ces*-positive or ces-negative state set to 0.2 and 0.8, respectively. Branch color corresponds to probability of a lineage being in a *ces*-negative state. Pie charts at nodes denote the posterior probability (PP) of a node being in a *ces*-negative (teal) or *ces*-positive (pink) state. Branch length is reported in substitutions/site/year. Core SNPs were identified using Snippy version 4.3.6. The phylogenies were constructed using the results of five independent runs using a relaxed lognormal clock model, the Standard_TVMef nucleotide substitution model, and the Birth Death Skyline Serial population model implemented in BEAST version 2.5.1, with 10% burn-in applied to each run. LogCombiner-2 was used to combine BEAST2 log files, and TreeAnnotator-2 was used to construct the phylogeny using common ancestor node heights.

**Supplemental Table S1.** Genomic data and metadata used in this study (*n* = 150).

**Supplemental Table S2.** Results of cereulide synthetase ancestral state reconstruction.

**Supplemental Text.** Detailed descriptions of all methods, plus references.

## References

1. Stenfors Arnesen LP, Fagerlund A, Granum PE. 2008. From soil to gut: *Bacillus cereus* and its food poisoning toxins. FEMS Microbiol Rev 32:579–606.

2. Kirk MD, Pires SM, Black RE, Caipo M, Crump JA, Devleesschauwer B, Dopfer D, Fazil A, Fischer-Walker CL, Hald T, Hall AJ, Keddy KH, Lake RJ, Lanata CF, Torgerson PR, Havelaar AH, Angulo FJ. 2015. World Health Organization Estimates of the Global and Regional Disease Burden of 22 Foodborne Bacterial, Protozoal, and Viral Diseases, 2010: A Data Synthesis. PLoS Med 12:e1001921.

3. Ehling-Schulz M, Fricker M, Scherer S. 2004. *Bacillus cereus*, the causative agent of an emetic type of food-borne illness. Mol Nutr Food Res 48:479–87.

4. Rajkovic A, Uyttendaele M, Vermeulen A, Andjelkovic M, Fitz-James I, in’t Veld P, Denon Q, Verhe R, Debevere J. 2008. Heat resistance of *Bacillus cereus* emetic toxin, cereulide. Lett Appl Microbiol 46:536–41.

5. Messelhäußer U, Ehling-Schulz M. 2018. *Bacillus cereus*—a Multifaceted Opportunistic Pathogen. Current Clinical Microbiology Reports 5:120–125.

6. Schoeni JL, Wong AC. 2005. *Bacillus cereus* food poisoning and its toxins. J Food Prot 68:636–48.

7. Carroll LM, Wiedmann M, Mukherjee M, Nicholas DC, Mingle LA, Dumas NB, Cole JA, Kovac J. 2019. Characterization of Emetic and Diarrheal *Bacillus cereus* Strains From a 2016 Foodborne Outbreak Using Whole-Genome Sequencing: Addressing the Microbiological, Epidemiological, and Bioinformatic Challenges. Front Microbiol 10:144.

8. Glasset B, Herbin S, Guillier L, Cadel-Six S, Vignaud ML, Grout J, Pairaud S, Michel V, Hennekinne JA, Ramarao N, Brisabois A. 2016. *Bacillus cereus*-induced food-borne outbreaks in France, 2007 to 2014: epidemiology and genetic characterisation. Euro Surveill 21.

9. Ehling-Schulz M, Fricker M, Grallert H, Rieck P, Wagner M, Scherer S. 2006. Cereulide synthetase gene cluster from emetic *Bacillus cereus*: structure and location on a mega virulence plasmid related to *Bacillus anthracis* toxin plasmid pXO1. BMC Microbiol 6:20.

10. Ehling-Schulz M, Frenzel E, Gohar M. 2015. Food-bacteria interplay: pathometabolism of emetic *Bacillus cereus*. Front Microbiol 6:704.

11. Guinebretiere MH, Thompson FL, Sorokin A, Normand P, Dawyndt P, Ehling-Schulz M, Svensson B, Sanchis V, Nguyen-The C, Heyndrickx M, De Vos P. 2008. Ecological diversification in the *Bacillus cereus* Group. Environ Microbiol 10:851–65.

12. Guinebretiere MH, Velge P, Couvert O, Carlin F, Debuyser ML, Nguyen-The C. 2010. Ability of *Bacillus cereus* group strains to cause food poisoning varies according to phylogenetic affiliation (groups I to VII) rather than species affiliation. J Clin Microbiol 48:3388–91.

13. Carroll LM, Kovac J, Miller RA, Wiedmann M. 2017. Rapid, high-throughput identification of anthrax-causing and emetic *Bacillus cereus* group genome assemblies using BTyper, a computational tool for virulence-based classification of *Bacillus cereus* group isolates using nucleotide sequencing data. Appl Environ Microbiol doi:10.1128/AEM.01096-17.

14. Hoton FM, Fornelos N, N’Guessan E, Hu X, Swiecicka I, Dierick K, Jaaskelainen E, Salkinoja-Salonen M, Mahillon J. 2009. Family portrait of *Bacillus cereus* and *Bacillus weihenstephanensis* cereulide-producing strains. Environ Microbiol Rep 1:177–83.

15. Guerin A, Ronning HT, Dargaignaratz C, Clavel T, Broussolle V, Mahillon J, Granum PE, Nguyen-The C. 2017. Cereulide production by *Bacillus weihenstephanensis* strains during growth at different pH values and temperatures. Food Microbiol 65:130–135.

16. Ehling-Schulz M, Svensson B, Guinebretiere MH, Lindback T, Andersson M, Schulz A, Fricker M, Christiansson A, Granum PE, Martlbauer E, Nguyen-The C, Salkinoja-Salonen M, Scherer S. 2005. Emetic toxin formation of *Bacillus cereus* is restricted to a single evolutionary lineage of closely related strains. Microbiology 151:183–197.

17. Thorsen L, Hansen BM, Nielsen KF, Hendriksen NB, Phipps RK, Budde BB. 2006. Characterization of emetic *Bacillus weihenstephanensis*, a new cereulide-producing bacterium. Appl Environ Microbiol 72:5118–21.

18. Castiaux V, N’Guessan E, Swiecicka I, Delbrassinne L, Dierick K, Mahillon J. 2014. Diversity of pulsed-field gel electrophoresis patterns of cereulide-producing isolates of *Bacillus cereus* and *Bacillus weihenstephanensis*. FEMS Microbiol Lett 353:124–31.

19. Mei X, Xu K, Yang L, Yuan Z, Mahillon J, Hu X. 2014. The genetic diversity of cereulide biosynthesis gene cluster indicates a composite transposon Tnces in emetic *Bacillus weihenstephanensis*. BMC Microbiol 14:149.

20. Tewari A, Abdullah S. 2015. *Bacillus cereus* food poisoning: international and Indian perspective. J Food Sci Technol 52:2500–11.

21. Carroll LM, Wiedmann M, Kovac J. 2020. Proposal of a Taxonomic Nomenclature for the *Bacillus cereus* Group Which Reconciles Genomic Definitions of Bacterial Species with Clinical and Industrial Phenotypes. mBio 11.

22. Arevalo P, VanInsberghe D, Elsherbini J, Gore J, Polz MF. 2019. A Reverse Ecology Approach Based on a Biological Definition of Microbial Populations. Cell 178:820–834 e14.

23. Miller RA, Beno SM, Kent DJ, Carroll LM, Martin NH, Boor KJ, Kovac J. 2016. *Bacillus wiedmannii* sp. nov., a psychrotolerant and cytotoxic *Bacillus cereus* group species isolated from dairy foods and dairy environments. Int J Syst Evol Microbiol 66:4744–4753.

24. Jimenez G, Urdiain M, Cifuentes A, Lopez-Lopez A, Blanch AR, Tamames J, Kampfer P, Kolsto AB, Ramon D, Martinez JF, Codoner FM, Rossello-Mora R. 2013. Description of *Bacillus toyonensis* sp. nov., a novel species of the *Bacillus cereus* group, and pairwise genome comparisons of the species of the group by means of ANI calculations. Syst Appl Microbiol 36:383–91.

25. Guinebretiere MH, Auger S, Galleron N, Contzen M, De Sarrau B, De Buyser ML, Lamberet G, Fagerlund A, Granum PE, Lereclus D, De Vos P, Nguyen-The C, Sorokin A. 2013. *Bacillus cytotoxicus* sp. nov. is a novel thermotolerant species of the *Bacillus cereus* Group occasionally associated with food poisoning. Int J Syst Evol Microbiol 63:31–40.

26. Liu Y, Du J, Lai Q, Zeng R, Ye D, Xu J, Shao Z. 2017. Proposal of nine novel species of the *Bacillus cereus* group. Int J Syst Evol Microbiol 67:2499–2508.

27. Naranjo M, Denayer S, Botteldoorn N, Delbrassinne L, Veys J, Waegenaere J, Sirtaine N, Driesen RB, Sipido KR, Mahillon J, Dierick K. 2011. Sudden death of a young adult associated with *Bacillus cereus* food poisoning. J Clin Microbiol 49:4379–81.

28. Dierick K, Van Coillie E, Swiecicka I, Meyfroidt G, Devlieger H, Meulemans A, Hoedemaekers G, Fourie L, Heyndrickx M, Mahillon J. 2005. Fatal family outbreak of *Bacillus cereus*-associated food poisoning. J Clin Microbiol 43:4277–9.

29. Mahler H, Pasi A, Kramer JM, Schulte P, Scoging AC, Bar W, Krahenbuhl S. 1997. Fulminant liver failure in association with the emetic toxin of *Bacillus cereus*. N Engl J Med 336:1142–8.

30. Posfay-Barbe KM, Schrenzel J, Frey J, Studer R, Korff C, Belli DC, Parvex P, Rimensberger PC, Schappi MG. 2008. Food poisoning as a cause of acute liver failure. Pediatr Infect Dis J 27:846–7.

31. Shiota M, Saitou K, Mizumoto H, Matsusaka M, Agata N, Nakayama M, Kage M, Tatsumi S, Okamoto A, Yamaguchi S, Ohta M, Hata D. 2010. Rapid detoxification of cereulide in *Bacillus cereus* food poisoning. Pediatrics 125:e951–5.

32. Yang Y, Gu H, Yu X, Zhan L, Chen J, Luo Y, Zhang Y, Zhang Y, Lu Y, Jiang J, Mei L. 2017. Genotypic heterogeneity of emetic toxin producing *Bacillus cereus* isolates from China. FEMS Microbiol Lett 364.

33. Vassileva M, Torii K, Oshimoto M, Okamoto A, Agata N, Yamada K, Hasegawa T, Ohta M. 2007. A new phylogenetic cluster of cereulide-producing *Bacillus cereus* strains. J Clin Microbiol 45:1274–7.

34. Apetroaie C, Andersson MA, Sproer C, Tsitko I, Shaheen R, Jaaskelainen EL, Wijnands LM, Heikkila R, Salkinoja-Salonen MS. 2005. Cereulide-producing strains of *Bacillus cereus* show diversity. Arch Microbiol 184:141–51.

35. Jain C, Rodriguez RL, Phillippy AM, Konstantinidis KT, Aluru S. 2018. High throughput ANI analysis of 90K prokaryotic genomes reveals clear species boundaries. Nat Commun 9:5114.

36. Worley J, Meng J, Allard MW, Brown EW, Timme RE. 2018. *Salmonella enterica* Phylogeny Based on Whole-Genome Sequencing Reveals Two New Clades and Novel Patterns of Horizontally Acquired Genetic Elements. MBio 9.

37. Usongo V, Berry C, Yousfi K, Doualla-Bell F, Labbe G, Johnson R, Fournier E, Nadon C, Goodridge L, Bekal S. 2018. Impact of the choice of reference genome on the ability of the core genome SNV methodology to distinguish strains of *Salmonella enterica* serovar Heidelberg. PLoS One 13:e0192233.

38. Olson ND, Lund SP, Colman RE, Foster JT, Sahl JW, Schupp JM, Keim P, Morrow JB, Salit ML, Zook JM. 2015. Best practices for evaluating single nucleotide variant calling methods for microbial genomics. Front Genet 6:235.

39. Pightling AW, Petronella N, Pagotto F. 2014. Choice of reference sequence and assembler for alignment of *Listeria monocytogenes* short-read sequence data greatly influences rates of error in SNP analyses. PLoS One 9:e104579.

40. Pightling AW, Petronella N, Pagotto F. 2015. Choice of reference-guided sequence assembler and SNP caller for analysis of *Listeria monocytogenes* short-read sequence data greatly influences rates of error. BMC Res Notes 8:748.

41. Hauge S. 1955. FOOD POISONING CAUSED BY AEROBIC SPORE-FORMING BACILLI. Journal of Applied Bacteriology 18:591–595.

42. Priest FG, Barker M, Baillie LW, Holmes EC, Maiden MC. 2004. Population structure and evolution of the *Bacillus cereus* group. J Bacteriol 186:7959–70.

43. Hoffmaster AR, Novak RT, Marston CK, Gee JE, Helsel L, Pruckler JM, Wilkins PP. 2008. Genetic diversity of clinical isolates of *Bacillus cereus* using multilocus sequence typing. BMC Microbiol 8:191.

44. Granum PE, Lund T. 1997. *Bacillus cereus* and its food poisoning toxins. FEMS Microbiol Lett 157:223–8.

45. Jessberger N, Krey VM, Rademacher C, Bohm ME, Mohr AK, Ehling-Schulz M, Scherer S, Martlbauer E. 2015. From genome to toxicity: a combinatory approach highlights the complexity of enterotoxin production in *Bacillus cereus*. Front Microbiol 6:560.

46. Riol CD, Dietrich R, Martlbauer E, Jessberger N. 2018. Consumed Foodstuffs Have a Crucial Impact on the Toxic Activity of Enteropathogenic *Bacillus cereus*. Front Microbiol 9:1946.

47. Hoton FM, Andrup L, Swiecicka I, Mahillon J. 2005. The cereulide genetic determinants of emetic *Bacillus cereus* are plasmid-borne. Microbiology 151:2121–2124.

48. Gdoura-Ben Amor M, Siala M, Zayani M, Grosset N, Smaoui S, Messadi-Akrout F, Baron F, Jan S, Gautier M, Gdoura R. 2018. Isolation, Identification, Prevalence, and Genetic Diversity of *Bacillus cereus* Group Bacteria From Different Foodstuffs in Tunisia. Front Microbiol 9:447.

49. Zhuang K, Li H, Zhang Z, Wu S, Zhang Y, Fox EM, Man C, Jiang Y. 2019. Typing and evaluating heat resistance of *Bacillus cereus sensu stricto* isolated from the processing environment of powdered infant formula. J Dairy Sci 102:7781–7793.

50. Glasset B, Herbin S, Granier SA, Cavalie L, Lafeuille E, Guerin C, Ruimy R, Casagrande-Magne F, Levast M, Chautemps N, Decousser JW, Belotti L, Pelloux I, Robert J, Brisabois A, Ramarao N. 2018. *Bacillus cereus*, a serious cause of nosocomial infections: Epidemiologic and genetic survey. PLoS One 13:e0194346.

51. Bukharin OV, Perunova NB, Andryuschenko SV, Ivanova EV, Bondarenko TA, Chainikova IN. 2019. Genome Sequence Announcement of *Bacillus paranthracis* Strain ICIS-279, Isolated from Human Intestine. Microbiol Resour Announc 8.

52. Pruitt KD, Tatusova T, Maglott DR. 2007. NCBI reference sequences (RefSeq): a curated non-redundant sequence database of genomes, transcripts and proteins. Nucleic Acids Res 35:D61–5.

53. Lechner S, Mayr R, Francis KP, Pruss BM, Kaplan T, Wiessner-Gunkel E, Stewart GS, Scherer S. 1998. *Bacillus weihenstephanensis* sp. nov. is a new psychrotolerant species of the *Bacillus cereus* group. Int J Syst Bacteriol 48 Pt 4:1373–82.

54. Jolley KA, Maiden MC. 2010. BIGSdb: Scalable analysis of bacterial genome variation at the population level. BMC Bioinformatics 11:595.

55. Gardner SN, Hall BG. 2013. When whole-genome alignments just won’t work: kSNP v2 software for alignment-free SNP discovery and phylogenetics of hundreds of microbial genomes. PLoS One 8:e81760.

56. Gardner SN, Slezak T, Hall BG. 2015. kSNP3.0: SNP detection and phylogenetic analysis of genomes without genome alignment or reference genome. Bioinformatics 31:2877–8.

57. Nguyen LT, Schmidt HA, von Haeseler A, Minh BQ. 2015. IQ-TREE: a fast and effective stochastic algorithm for estimating maximum-likelihood phylogenies. Mol Biol Evol 32:268–74.

58. Kalyaanamoorthy S, Minh BQ, Wong TKF, von Haeseler A, Jermiin LS. 2017. ModelFinder: fast model selection for accurate phylogenetic estimates. Nat Methods 14:587–589.

59. Minh BQ, Nguyen MA, von Haeseler A. 2013. Ultrafast approximation for phylogenetic bootstrap. Mol Biol Evol 30:1188–95.

60. Hoang DT, Chernomor O, von Haeseler A, Minh BQ, Vinh LS. 2018. UFBoot2: Improving the Ultrafast Bootstrap Approximation. Mol Biol Evol 35:518–522.

61. Revell LJ. 2012. phytools: an R package for phylogenetic comparative biology (and other things). Methods in Ecology and Evolution 3:217–223.

62. Tonkin-Hill G, Lees JA, Bentley SD, Frost SDW, Corander J. 2018. RhierBAPS: An R implementation of the population clustering algorithm hierBAPS. Wellcome Open Res 3:93.

63. Seemann T. 2019. Snippy: Rapid haploid variant calling and core genome alignment, v4.3.6. https://github.com/tseemann/snippy.

64. Croucher NJ, Page AJ, Connor TR, Delaney AJ, Keane JA, Bentley SD, Parkhill J, Harris SR. 2015. Rapid phylogenetic analysis of large samples of recombinant bacterial whole genome sequences using Gubbins. Nucleic Acids Res 43:e15.

65. Page AJ, Taylor B, Delaney AJ, Soares J, Seemann T, Keane JA, Harris SR. 2016. SNP-sites: rapid efficient extraction of SNPs from multi-FASTA alignments. Microb Genom 2:e000056.

66. Rambaut A, Lam TT, Max Carvalho L, Pybus OG. 2016. Exploring the temporal structure of heterochronous sequences using TempEst (formerly Path-O-Gen). Virus Evol 2:vew007.

67. Bouckaert R, Vaughan TG, Barido-Sottani J, Duchene S, Fourment M, Gavryushkina A, Heled J, Jones G, Kuhnert D, De Maio N, Matschiner M, Mendes FK, Muller NF, Ogilvie HA, du Plessis L, Popinga A, Rambaut A, Rasmussen D, Siveroni I, Suchard MA, Wu CH, Xie D, Zhang C, Stadler T, Drummond AJ. 2019. BEAST 2.5: An advanced software platform for Bayesian evolutionary analysis. PLoS Comput Biol 15:e1006650.

68. Bouckaert R, Heled J, Kuhnert D, Vaughan T, Wu CH, Xie D, Suchard MA, Rambaut A, Drummond AJ. 2014. BEAST 2: a software platform for Bayesian evolutionary analysis. PLoS Comput Biol 10:e1003537.

69. Bouckaert R, Xie D. 2017. SSN: Standard Nucleotide Substitution Models, http://doi.org/10.5281/zenodo.995740.

70. Bouckaert R. 2014. Correcting for constant sites in BEAST2. https://groups.google.com/forum/#!topic/beast-users/QfBHMOqImFE. Accessed May 12, 2020.

71. Drummond AJ, Ho SY, Phillips MJ, Rambaut A. 2006. Relaxed phylogenetics and dating with confidence. PLoS Biol 4:e88.

72. Stadler T, Kuhnert D, Bonhoeffer S, Drummond AJ. 2013. Birth-death skyline plot reveals temporal changes of epidemic spread in HIV and hepatitis C virus (HCV). Proc Natl Acad Sci U S A 110:228–33.

73. Rambaut A, Drummond AJ, Xie D, Baele G, Suchard MA. 2018. Posterior Summarization in Bayesian Phylogenetics Using Tracer 1.7. Syst Biol 67:901–904.

74. Heled J, Bouckaert RR. 2013. Looking for trees in the forest: summary tree from posterior samples. BMC Evol Biol 13:221.

75. Yoshimura D, Kajitani R, Gotoh Y, Katahira K, Okuno M, Ogura Y, Hayashi T, Itoh T. 2019. Evaluation of SNP calling methods for closely related bacterial isolates and a novel high-accuracy pipeline: BactSNP. Microb Genom 5.

76. Katz LS, Griswold T, Williams-Newkirk AJ, Wagner D, Petkau A, Sieffert C, Van Domselaar G, Deng X, Carleton HA. 2017. A Comparative Analysis of the Lyve-SET Phylogenomics Pipeline for Genomic Epidemiology of Foodborne Pathogens. Front Microbiol 8:375.

77. Treangen TJ, Ondov BD, Koren S, Phillippy AM. 2014. The Harvest suite for rapid core-genome alignment and visualization of thousands of intraspecific microbial genomes. Genome Biol 15:524.

78. Bruen TC, Philippe H, Bryant D. 2006. A simple and robust statistical test for detecting the presence of recombination. Genetics 172:2665–81.

79. Kendall M, Colijn C. 2016. Mapping Phylogenetic Trees to Reveal Distinct Patterns of Evolution. Molecular Biology and Evolution 33:2735–2743.

80. Kendall M, Colijn C. 2015. A tree metric using structure and length to capture distinct phylogenetic signals. arXiv:1507.05211.

81. Zwick ME, Joseph SJ, Didelot X, Chen PE, Bishop-Lilly KA, Stewart AC, Willner K, Nolan N, Lentz S, Thomason MK, Sozhamannan S, Mateczun AJ, Du L, Read TD. 2012. Genomic characterization of the *Bacillus cereus sensu lato* species: backdrop to the evolution of *Bacillus anthracis*. Genome Res 22:1512–24.

82. Xiong Z, Jiang Y, Qi D, Lu H, Yang F, Yang J, Chen L, Sun L, Xu X, Xue Y, Zhu Y, Jin Q. 2009. Complete genome sequence of the extremophilic *Bacillus cereus* strain Q1 with industrial applications. J Bacteriol 191:1120–1.

83. Ji F, Zhu Y, Ju S, Zhang R, Yu Z, Sun M. 2009. Promoters of crystal protein genes do not control crystal formation inside exosporium of *Bacillus thuringiensis* ssp. *finitimus* strain YBT-020. FEMS Microbiol Lett 300:11–7.

84. Guo G, Zhang L, Zhou Z, Ma Q, Liu J, Zhu C, Zhu L, Yu Z, Sun M. 2008. A new group of parasporal inclusions encoded by the S-layer gene of *Bacillus thuringiensis*. FEMS Microbiol Lett 282:1–7.

85. Zhu Y, Ji F, Shang H, Zhu Q, Wang P, Xu C, Deng Y, Peng D, Ruan L, Sun M. 2011. Gene clusters located on two large plasmids determine spore crystal association (SCA) in *Bacillus thuringiensis* subsp. *finitimus* strain YBT-020. PLoS One 6:e27164.

86. Zhu Y, Shang H, Zhu Q, Ji F, Wang P, Fu J, Deng Y, Xu C, Ye W, Zheng J, Zhu L, Ruan L, Peng D, Sun M. 2011. Complete genome sequence of *Bacillus thuringiensis* serovar *finitimus* strain YBT-020. J Bacteriol 193:2379–80.

87. Fiedoruk K, Daniluk T, Fiodor A, Drewicka E, Buczynska K, Leszczynska K, Bideshi DK, Swiecicka I. 2016. MALDI-TOF MS portrait of emetic and non-emetic *Bacillus cereus* group members. Electrophoresis 37:2235–47.

88. Su L, Zhou T, Zhou L, Fang X, Li T, Wang J, Guo Y, Chang D, Wang Y, Li D, Liu C. 2012. Draft genome sequence of *Bacillus cereus* strain LCT-BC244. J Bacteriol 194:3549.

89. Agata N, Mori M, Ohta M, Suwan S, Ohtani I, Isobe M. 1994. A novel dodecadepsipeptide, cereulide, isolated from *Bacillus cereus* causes vacuole formation in HEp-2 cells. FEMS Microbiol Lett 121:31–4.

90. Ekman JV, Kruglov A, Andersson MA, Mikkola R, Raulio M, Salkinoja-Salonen M. 2012. Cereulide produced by *Bacillus cereus* increases the fitness of the producer organism in low-potassium environments. Microbiology 158:1106–1116.

91. Takeno A, Okamoto A, Tori K, Oshima K, Hirakawa H, Toh H, Agata N, Yamada K, Ogasawara N, Hayashi T, Shimizu T, Kuhara S, Hattori M, Ohta M. 2012. Complete genome sequence of *Bacillus cereus* NC7401, which produces high levels of the emetic toxin cereulide. J Bacteriol 194:4767–8.

92. Hu X, Van der Auwera G, Timmery S, Zhu L, Mahillon J. 2009. Distribution, diversity, and potential mobility of extrachromosomal elements related to the *Bacillus anthracis* pXO1 and pXO2 virulence plasmids. Appl Environ Microbiol 75:3016–28.

93. Van der Auwera GA, Feldgarden M, Kolter R, Mahillon J. 2013. Whole-Genome Sequences of 94 Environmental Isolates of *Bacillus cereus Sensu Lato*. Genome Announc 1.

94. Swiecicka I, De Vos P. 2003. Properties of *Bacillus thuringiensis* isolated from bank voles. J Appl Microbiol 94:60–4.

95. Biodefense and Emerging Infections (BEI) Research Resources Repository. 2019. Bacillus cereus Strain AND1407, NR-22159. https://www.beiresources.org/Catalog/Bacteria/NR-22159.aspx. Accessed December 24, 2019.

96. Timmery S, Hu X, Mahillon J. 2011. Characterization of Bacilli isolated from the confined environments of the Antarctic Concordia station and the International Space Station. Astrobiology 11:323–34.

97. Zhang X, Wang T, Su L, Zhou L, Li T, Wang J, Liu Y, Jiang X, Wu C, Liu C. 2014. Draft Genome Sequence of *Bacillus cereus* LCT-BC25, Isolated from Space Flight. Genome Announc 2.

98. Su L, Wang T, Zhou L, Wu C, Guo Y, Chang D, Liu Y, Jiang X, Yin S, Liu C. 2014. Genome Sequence of *Bacillus cereus* Strain LCT-BC235, Carried by the Shenzhou VIII Spacecraft. Genome Announc 2.

99. Radnedge L, Agron PG, Hill KK, Jackson PJ, Ticknor LO, Keim P, Andersen GL. 2003. Genome differences that distinguish *Bacillus anthracis* from *Bacillus cereus* and *Bacillus thuringiensis*. Appl Environ Microbiol 69:2755–64.

100. Zhong W, Shou Y, Yoshida TM, Marrone BL. 2007. Differentiation of *Bacillus anthracis*, *B. cereus*, and *B. thuringiensis* by using pulsed-field gel electrophoresis. Appl Environ Microbiol 73:3446–9.

101. Pannucci J, Okinaka RT, Sabin R, Kuske CR. 2002. *Bacillus anthracis* pXO1 plasmid sequence conservation among closely related bacterial species. J Bacteriol 184:134–41.

102. Knight BC, Proom H. 1950. A comparative survey of the nutrition and physiology of mesophilic species in the genus *Bacillus*. J Gen Microbiol 4:508–38.

103. Sneath PH. 1955. Proof of the spontaneity of a mutation to penicillinase production in *Bacillus cereus*. J Gen Microbiol 13:561–8.

104. Fenselau C, Havey C, Teerakulkittipong N, Swatkoski S, Laine O, Edwards N. 2008. Identification of beta-lactamase in antibiotic-resistant *Bacillus cereus* spores. Appl Environ Microbiol 74:904–6.

105. Krawczyk AO, de Jong A, Eijlander RT, Berendsen EM, Holsappel S, Wells-Bennik MH, Kuipers OP. 2015. Next-Generation Whole-Genome Sequencing of Eight Strains of *Bacillus cereus*, Isolated from Food. Genome Announc 3.

106. Bohm ME, Huptas C, Krey VM, Scherer S. 2015. Massive horizontal gene transfer, strictly vertical inheritance and ancient duplications differentially shape the evolution of *Bacillus cereus* enterotoxin operons *hbl*, *cytK* and *nhe*. BMC Evol Biol 15:246.

107. Crovadore J, Calmin G, Tonacini J, Chablais R, Schnyder B, Messelhausser U, Lefort F. 2016. Whole-Genome Sequences of Seven Strains of *Bacillus cereus* Isolated from Foodstuff or Poisoning Incidents. Genome Announc 4.

108. Miller RA, Jian J, Beno SM, Wiedmann M, Kovac J. 2018. Intraclade Variability in Toxin Production and Cytotoxicity of *Bacillus cereus* Group Type Strains and Dairy-Associated Isolates. Appl Environ Microbiol 84.

109. Kovac J, Miller RA, Carroll LM, Kent DJ, Jian J, Beno SM, Wiedmann M. 2016. Production of hemolysin BL by *Bacillus cereus* group isolates of dairy origin is associated with whole-genome phylogenetic clade. BMC Genomics 17:581.

110. Hayrapetyan H, Boekhorst J, de Jong A, Kuipers OP, Nierop Groot MN, Abee T. 2016. Draft Whole-Genome Sequences of 11 *Bacillus cereus* Food Isolates. Genome Announc 4.

111. Zeigler DR. 1999. *Bacillus* Genetic Stock Center Catalog of Strains, Seventh Edition, Part 2: *Bacillus thuringiensis* and *Bacillus cereus* 7th ed. The Bacillus Genetic Stock Center, The Ohio State University, Columbus, Ohio.

112. Raymond B. 2017. The Biology, Ecology and Taxonomy of *Bacillus thuringiensis* and Related Bacteria, p 19–39. *In* Fiuza LM, Polanczyk RA, Crickmore N (ed), Bacillus thuringiensis and Lysinibacillus sphaericus: Characterization and use in the field of biocontrol doi:10.1007/978-3-319-56678-8_2. Springer International Publishing, Cham.

113. Liu M, Cai QX, Liu HZ, Zhang BH, Yan JP, Yuan ZM. 2002. Chitinolytic activities in *Bacillus thuringiensis* and their synergistic effects on larvicidal activity. J Appl Microbiol 93:374–9.

114. Che L, Xu W, Zhan J, Zhang L, Liu L, Zhou H. 2019. Complete Genome Sequence of *Bacillus cereus* CC-1, A Novel Marine Selenate/Selenite Reducing Bacterium Producing Metallic Selenides Nanomaterials. Curr Microbiol 76:78–85.

115. Grubbs KJ, Bleich RM, Santa Maria KC, Allen SE, Farag S, AgBiome T, Shank EA, Bowers AA. 2017. Large-Scale Bioinformatics Analysis of *Bacillus* Genomes Uncovers Conserved Roles of Natural Products in Bacterial Physiology. mSystems 2.

116. Chang T, Rosch JW, Gu Z, Hakim H, Hewitt C, Gaur A, Wu G, Hayden RT. 2018. Whole-Genome Characterization of *Bacillus cereus* Associated with Specific Disease Manifestations. Infect Immun 86.

117. Shankar M, Mageswari A, Suganthi C, Gunasekaran P, Gothandam KM, Karthikeyan S. 2018. Genome Sequence of a Moderately Halophilic *Bacillus cereus* Strain, TS2, Isolated from Saltern Sediments. Microbiol Resour Announc 7.

118. Ikram S, Heikal A, Finke S, Hofgaard A, Rehman Y, Sabri AN, Okstad OA. 2019. *Bacillus cereus* biofilm formation on central venous catheters of hospitalised cardiac patients. Biofouling 35:204–216.

