## Supplemental Figure S2 for "Cereulide synthetase acquisition and loss events within the evolutionary history of Group III *Bacillus cereus sensu lato* facilitate the transition between emetic and diarrheal foodborne pathogen"

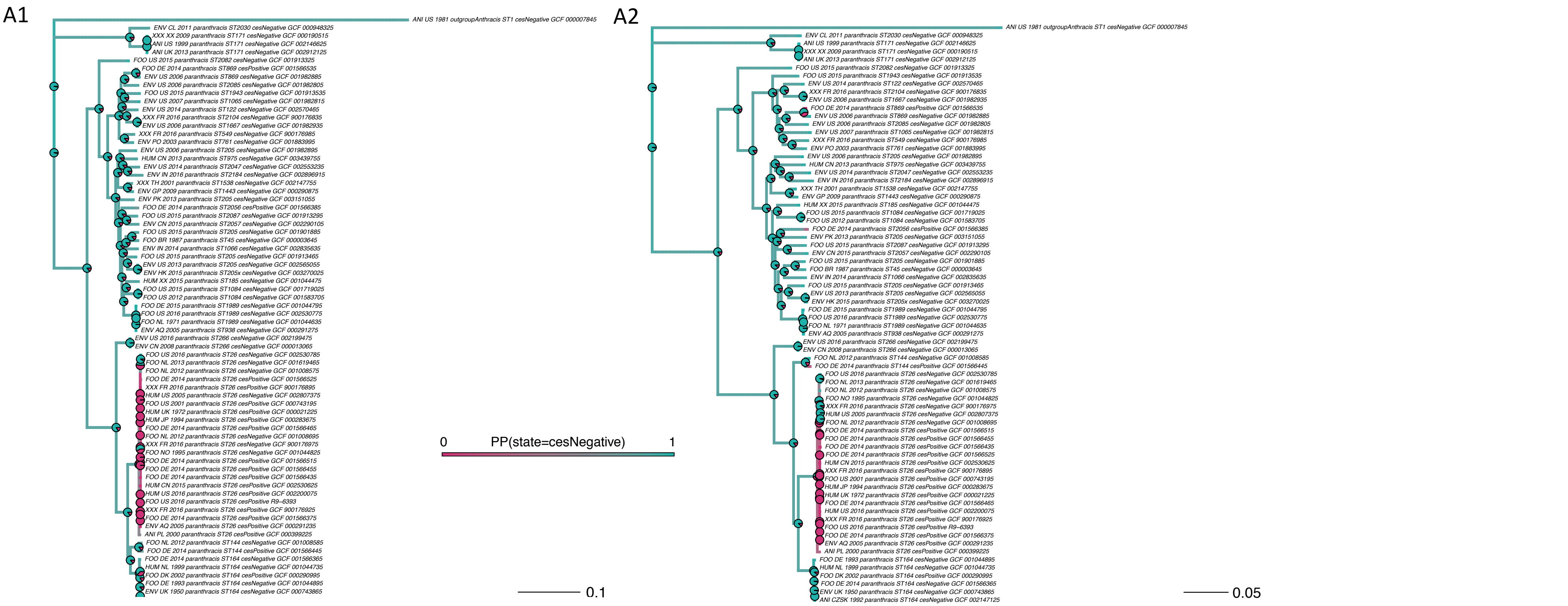


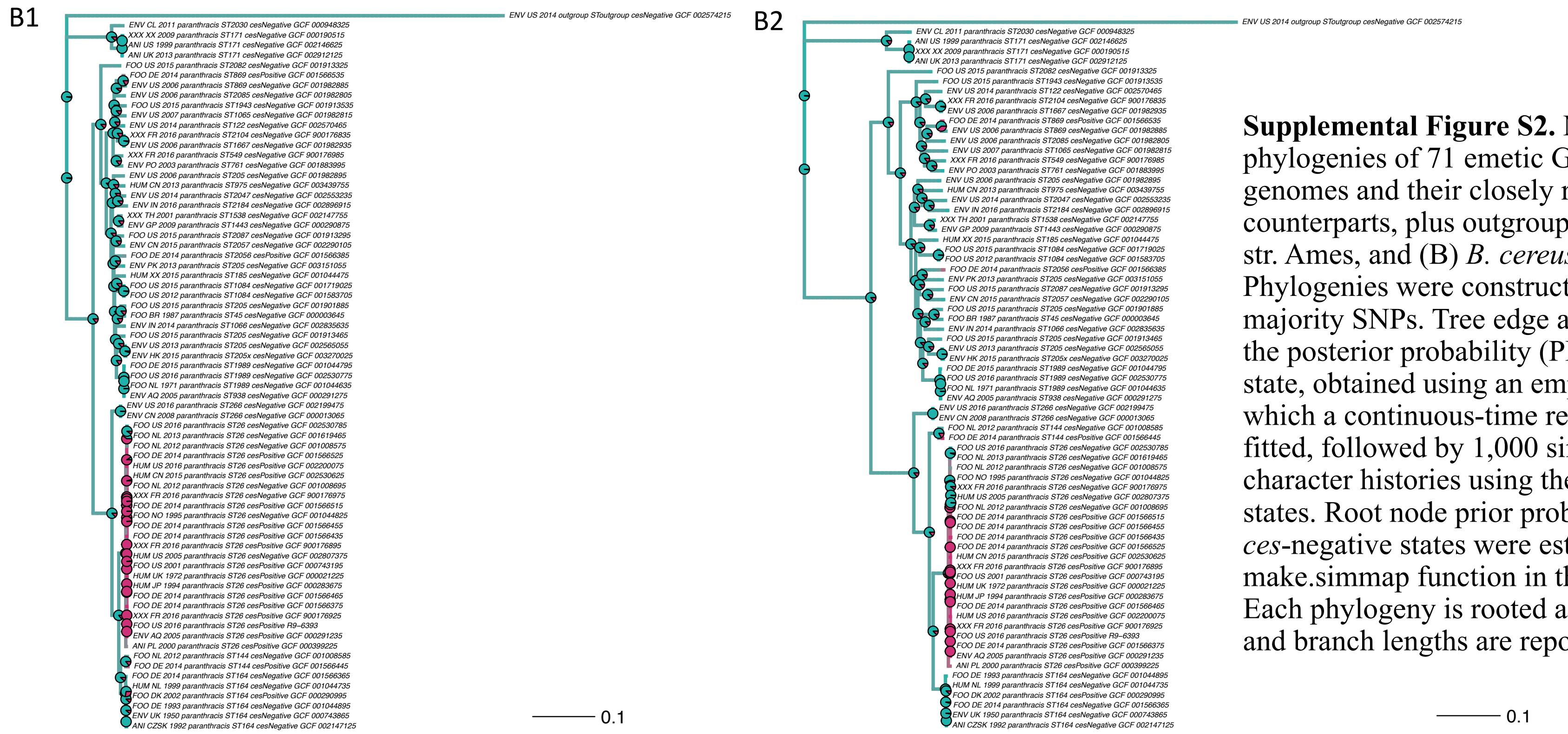


**Supplemental Figure S2.** Maximum likelihood phylogenies of 71 emetic Group III *B. cereus s.l.* genomes and their closely related, non-emetic counterparts, plus outgroup genomes (A) *B. anthracis* str. Ames, and (B) *B. cereus s.l.* str. AFS057383. Phylogenies were constructed using (1) core, and (2) majority SNPs. Tree edge and node colors correspond to the posterior probability (PP) of being in a *ces*-negative state, obtained using an empirical Bayes approach, in which a continuous-time reversible Markov model was fitted, followed by 1,000 simulations of stochastic character histories using the fitted model and tree tip states. Root node prior probabilities for *ces*-positive and *ces*-negative states were estimated using the make.simmap function in the phytools package in R. Each phylogeny is rooted along its respective outgroup, and branch lengths are reported in substitutions/site.
