## Supplemental Figure S3 for "Cereulide synthetase acquisition and loss events within the evolutionary history of Group III *Bacillus cereus sensu lato* facilitate the transition between emetic and diarrheal foodborne pathogen"

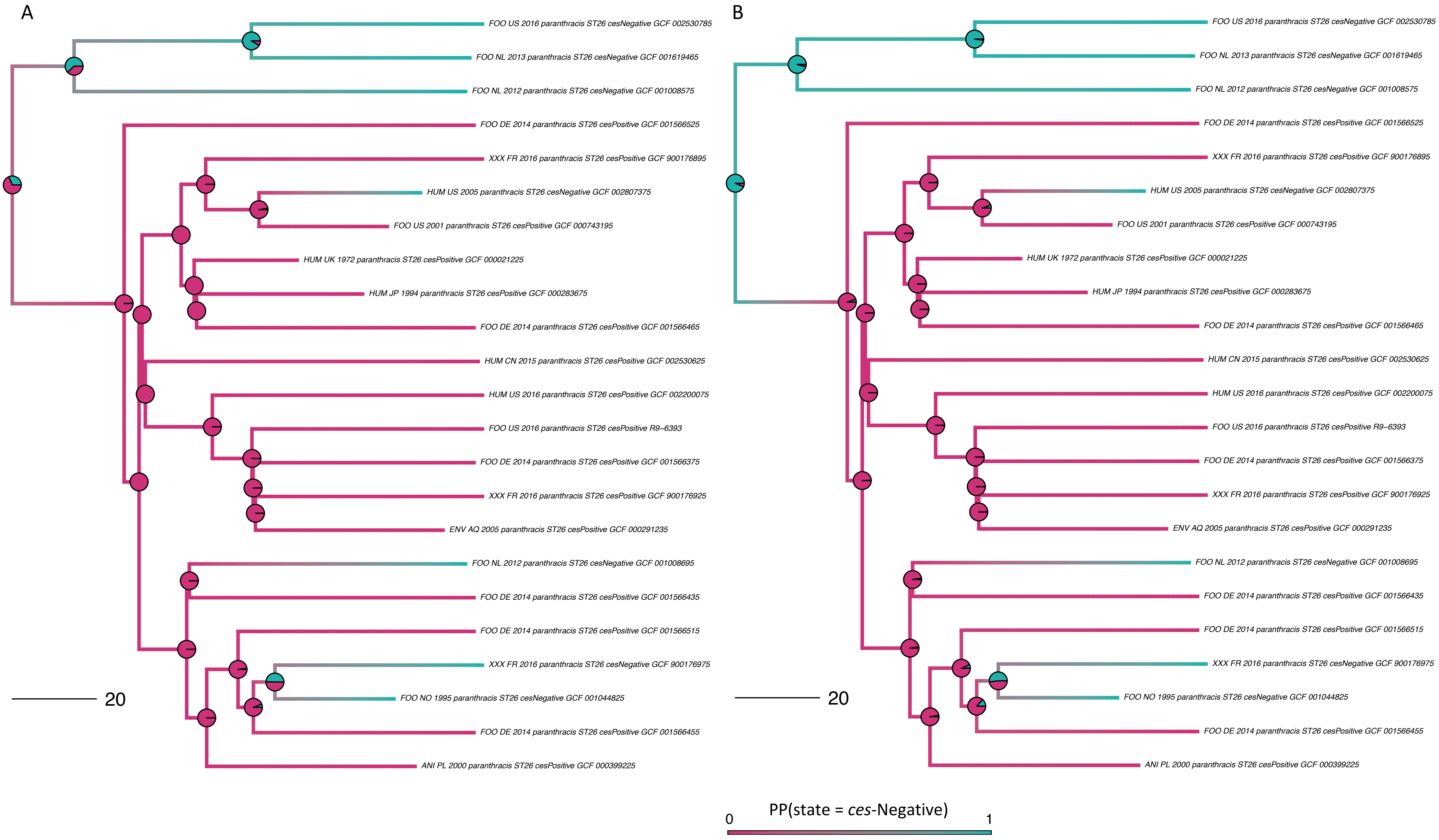


**Supplemental Figure S3.** Rooted, time-scaled maximum clade credibility (MCC) phylogenies constructed using core SNPs identified among 23 Group III *B. cereus s.l.* genomes belonging to sequence type (ST) 26. Ancestral state reconstruction was performed using the following priors on the root node: (A) probability of the root node belonging to a *ces*-positive or *ces*-negative state set to 0.5 each; or (B) probability of the root node being in a *ces*-positive or ces-negative state set to 0.2 and 0.8, respectively. Branch color corresponds to probability of a lineage being in a *ces*-negative state. Pie charts at nodes denote the posterior probability (PP) of a node being in a *ces*-negative (teal) or *ces*-positive (pink) state. Branch length is reported in substitutions/site/year. Core SNPs were identified using Snippy version 4.3.6. The phylogenies were constructed using the results of five independent runs using a relaxed lognormal clock model, the Standard_TVMef nucleotide substitution model, and the Birth Death Skyline Serial population model implemented in BEAST version 2.5.1, with 10% burn-in applied to each run. LogCombiner-2 was used to combine BEAST2 log files, and TreeAnnotator-2 was used to construct the phylogeny using common ancestor node heights.
