## Supplemental Text for "Cereulide synthetase acquisition and loss events within the evolutionary history of Group III *Bacillus cereus sensu lato* facilitate the transition between emetic and diarrheal foodborne pathogen"

Detailed descriptions of all methods, plus references.

**Acquisition of Group III *B. cereus s.l.* genomes and metadata.** All genomes submitted to the National Center for Biotechnology Information (NCBI) RefSeq (1) database under the name of a published species belonging to *B. cereus s.l.* (i.e., one of *B. albus, anthracis, cereus, cytotoxicus, luti, mobilis, mycoides, nitratireducens, pacificus, paramycoides, paranthracis, proteolyticus, pseudomycoides, thuringiensis, toyonensis, tropicus, weihenstephanensis,* or *wiedmannii*) (2-7) were downloaded ($n$ = 2,231; accessed November 19, 2018). The one-way average nucleotide identity BLAST (ANIb) function in BTyper version 2.3.3 (8) was used to calculate ANIb values between each of the 2,231 assembled *B. cereus s.l.* genomes and genomes of each of the 18 published *B. cereus s.l.* species as they existed in 2019 (for all but *B. anthracis*, the species type strain genome was used; for *B. anthracis*, the closed chromosome of *B. anthracis* str. Ames was used, as it is the reference genome for the species and the only type strain genome was scaffolded). *B. cereus s.l.* genomes which (i) most closely resembled the *B. paranthracis* type strain genome (i.e., the highest ANIb value was produced when the genome was compared to *B. paranthracis*), and (ii) shared an ANIb value ≥ 95 with the *B. paranthracis* type strain genome were used in subsequent steps ($n$ = 120), as this set of genomes contained all Group III *B. cereus s.l.* genomes that possessed genes encoding cereulide synthetase (described in detail below). The resulting 120 Group III *B. cereus s.l.* genomes were supplemented with an additional 30 Group III *B. cereus s.l.* genomes of strains isolated in conjunction with a 2016 emetic outbreak in New York State (9), resulting in a total of 150 Group III *B. cereus s.l.* genomes (Supplemental Table S1). FastANI version 1.0 (10) was used to confirm that all 150 genomes selected for this study (i) shared ≥ 95 ANI with the *B. paranthracis* type strain genome, and (ii) most closely resembled the *B. paranthracis* type strain genome when compared to the 18 *B. cereus s.l.* type strain/reference genomes.

Metadata for each of the 150 Group III *B. cereus s.l.* genomes was obtained using publicly available records. First, the NCBI BioSample (11) associated with each genome assembly was queried for (i) isolation source, (ii) geographic location, and (iii) year of isolation. If any of this information was not available within the BioSample record, the BioProject linked to the BioSample was queried. If this search did not return additional metadata, any publications (e.g., research papers, genome announcements) linked to the BioProject were queried. Finally, strain names of genomes without linked publications were queried in Google to obtain possible unlinked publications or hits in additional public databases. Using metadata that resulted from these searches, each genome was assigned (i) an isolation source, (ii) a geographic location, and (iii) a year of isolation. For isolation source, genomes were categorized into one of the following groups: ANI (isolated from an animal, excluding humans), ENV (isolated from an environment not meant for human consumption), FOO (isolated directly from a food product, food ingredient, or dietary supplement with the potential for human consumption), HUM (isolated from a human), and XXX (isolated from an unknown source) (Supplemental Table S1). For geographic location, isolates were grouped by their country of isolation, except for a few cases in which a major autonomous region was listed (i.e., Hong Kong), a country which no longer existed was listed (i.e., Czechoslovakia), or a country of isolation designation was not applicable (i.e., isolation occurred in the Pacific Ocean or Antarctica) (Supplemental Table S1). Isolates which could not be assigned a country of isolation were given a geographic isolation designation of XX. For year of isolation, genomes of strains with an "exact" year of isolation listed in a public database or publication were assigned to that particular year (Supplemental Table S1). For genomes for which this information was unavailable, a "maximum year of isolation" which corresponded to the year associated with the earliest appearance of the strain in a publication or public resource (e.g., database or strain collection) was assigned (Supplemental Table S1).

Each of the 150 Group III *B. cereus s.l.* genomes were additionally assigned a sequence type (ST), as well as a designation of potentially emetic or not. To assess the emetic potential of each of the 150 Group III *B. cereus s.l.* genomes, BTyper version 2.3.3 was used to detect cereulide synthetase genes *cesABCD* in each assembly, first using the default coverage and identity thresholds (70 and 50%, respectively), and a second time with 0% coverage to ensure that *cesABCD* were absent from genomes in which the genes were not detected (the only genome which was affected by this was that of one of the outbreak isolates, FSL R9-6384, which had *cesD* split on two contigs). All isolates in which any of *cesABCD* were detected possessed all four genes; these isolates were given a designation of *ces*-positive with the potential to cause emetic disease. Isolates in which *cesABCD* were not detected were given a designation of *ces*-negative. BTyper was additionally used to detect *cesABCD* in each of the 2,111 *B. cereus s.l.* genomes not included in this study, as well as to assign all *B. cereus s.l.* genomes to a *panC* group using the typing scheme described by Guinebretiere, et al (12). All 150 *B. cereus s.l.* genomes used in this study were assigned to *panC* Group III, and all Group III genomes possessing *cesABCD* were confirmed to have been included in this study. The only other genomes that possessed *cesABCD* belonged to *panC* group VI and most closely resembled the type strain genomes of *B. mycoides*/*B. weihenstephanensis* (referred to previously as "emetic *B. weihenstephanensis*") (7). BTyper version 2.3.3 was also used to assign each genome to a ST using the seven-gene multi-locus sequence typing (MLST) scheme available in PubMLST (13).

One genome (NCBI RefSeq Accession GCF_003270025) was assigned a probable ST of 205 but had mismatches in the *gmk* and *tpi* loci; as a result, a "x" character was appended after its ST to denote this (ST205x; Supplemental Table S1).

Using metadata and typing results obtained as described above, each genome was assigned a strain name adhering to the following format: (i) isolation source, (ii) geographic location, (iii) year of isolation, (iv) ANI-assigned species (i.e., *paranthracis*, the proposed species definition in use in 2018; note that a recently published taxonomic framework proposes the use of *mosaicus*) (7), (v) MLST-assigned ST, (vi) *ces*-positive or *ces*-negative designation, and (vii) RefSeq assembly accession or Food Microbe Tracker Strain identifier (14) for genomes obtained from RefSeq or the foodborne outbreak described by Carroll et al., respectively (Supplemental Table S1) (9). The rationale for assigning each genome a particular isolation source, geographic location, or isolation year can be found in Supplemental Table S1.

**Construction of Group III *B. cereus s.l.* maximum likelihood phylogenies and ancestral state reconstruction.** kSNP3 version 3.1 (15, 16) was used to identify SNPs among genomes in the following data sets: (i) all 150 Group III *B. cereus s.l.* genomes described above, plus the closed RefSeq species reference genome for *B. anthracis* (*B. anthracis* str. Ames, NCBI RefSeq Accession GCF_000007845.1; this genome would be treated as an outgroup for ancestral state reconstruction), and (ii) all 150 Group III *B. cereus s.l.* genomes described above, plus the draft genome of *B. cereus s.l.* strain AFS057383 (NCBI RefSeq Accession GCF_002574215.1; this genome would also be treated as an outgroup to ensure that choice of outgroup did not affect ancestral state reconstruction). For both data sets, Kchooser was used to determine the optimal *k*-mer size ($k = 21$ for both). Alignments of (i) core and (ii) majority (i.e., detected in > 50% of all genomes in the alignment) SNPs detected among the 150 Group III *B. cereus s.l.* genomes in this study, plus one of two outgroup genomes (i.e., either *B. anthracis* str. Ames or *B. cereus s.l.* str. AFS057383) using kSNP3 were used as input for IQ-TREE version 1.6.10 (17). For each of the four SNP alignments (i.e., each combination of outgroup and either core or majority SNPs), the optimal ascertainment bias-aware (18) nucleotide substitution model selected using ModelFinder (i.e., the model with the lowest Bayesian Information Criterion [BIC] value) was used (19), and branch support was assessed using 1,000 replicates of the ultrafast bootstrap approximation (20, 21).

To estimate ancestral character states of internal nodes in the Group III *B. cereus s.l.* phylogeny as they related to cereulide production (i.e., whether a node in the tree represents an ancestor that is more likely to be *ces*-positive or *ces*-negative), the presence or absence of *ces* within each genome was treated as a binary state. Each of the four phylogenies constructed as described above was rooted at its respective outgroup (i.e., either *B. anthracis* str. Ames or *B. cereus s.l.* str. AFS057383) using the root function in the ape package (22, 23) in R version 3.6.1 (24). Stochastic character maps were simulated on each of the four phylogenies using the make.simmap function in the phytools package (25) and the all-rates-different (ARD) model in the ape package. For each of the four phylogenies, either (i) equal root node prior probabilities for *ces*-positive and *ces*-negative states (i.e., $P(ces\ present) = 0.5$ and $P(ces\ absent) = 0.5$), or (ii) estimated root node prior probabilities for *ces*-positive and *ces*-negative states obtained using the make.simmap function were used. For each root node prior/phylogeny combination (eight total combinations of two root node priors and four phylogenies), an empirical Bayes approach was used, in which a continuous-time reversible Markov model was fitted, followed by

1,000 simulations of stochastic character histories using the fitted model and tree tip states (Supplemental Table S2). The resulting phylogenies were plotted using the densityMap function in the phytools package.

To ensure that ancestral state reconstruction would not be affected by genomes of isolates over-represented in RefSeq (e.g., genomes confirmed or predicted to have been derived from strains isolated from the same outbreak), potential duplicate genomes were removed using isolate metadata and by assessing isolate clustering in the ML phylogenies. One representative genome was selected from clusters that likely consisted of duplicate genomes and/or isolates derived from the same source. For example, this procedure reduced 30 closely related isolates from a 2016 outbreak (9) to one isolate. Overall, this approach yielded a reduced, de-replicated set of 71 Group III *B. cereus s.l.* genomes (Supplemental Table S1). kSNP and IQ-TREE were again used to identify core and majority SNPs and construct ML phylogenies among the set of 71 de-replicated genomes, plus each of the two outgroup genomes, and ancestral state reconstruction was performed as described above (for both data sets, the optimal *k*-mer size determined by Kchooser was 23).

**Assessment of Group III *B. cereus s.l.* population structure.** kSNP3 version 3.1 was used to identify core SNPs among the de-replicated set of 71 Group III *B. cereus s.l.* genomes, using the optimal *k*-mer size selected by Kchooser (*k* = 23). The set of core SNPs produced by kSNP3 was used as input for RhierBAPS (26) to identify clusters among the 71 genomes, using two clustering levels. The same set of 71 genomes was used as input for PopCOGenT (downloaded October 5, 2019) to identify gene flow among sub-populations of Group III *B. cereus s.l.* genomes (27), using Mugsy version v1r2.3 (28), PhyML version 20120412 patch 20131031 (29),

MMseqs2 version 67c04ae456664d910059dc194863451475d2e15a (30), and MUSCLE version 3.8.31 (31).

**Construction of Group III *B. cereus s.l.* ST 26 temporal phylogeny.** Snippy version 4.3.6 (32) was used to identify core SNPs among the de-replicated set of 23 ST 26 genomes (see section "Construction of Group III *B. cereus s.l.* maximum likelihood phylogenies and ancestral state reconstruction" above), using the closed chromosome of emetic *B. cereus s.l.* ST 26 str. AH187 (NCBI RefSeq Assession NC_011658.1) as a reference genome and the following software as dependencies: BWA MEM version 0.7.13-r1126 (33, 34), Minimap2 version 2.15 (35), SAMtools version 1.8 (36), BEDtools version 2.27.1 (37, 38), BCFtools version 1.8 (39), FreeBayes version v1.1.0-60-gc15b070 (40), vcflib version v1.0.0-rc2 (41), vt version 0.57721 (42), SnpEff version 4.3T (43), samclip version 0.2 (44), seqtk version 1.2-r102-dirty (45), and snp-sites version 2.4.0 (46). Three isolates had Illumina short reads of adequate quality after trimming and adapter removal using Trimmomatic 0.39 (47) (as determined using FastQC version 0.11.8) (48); as such, reads were used as input for these isolates, and assembled genomes were used for the remaining 20 ST 26 isolates. Gubbins version 2.3.4 (49) was used to remove recombination from the resulting alignment, and snp-sites was used to obtain core SNPs among the 23 genomes. IQ-TREE was used to construct a ML phylogeny, using the ST 26 core SNP alignment as input, the optimal ascertainment bias-aware nucleotide substitution model selected using ModelFinder, and 1,000 replicates of the ultrafast bootstrap approximation. The temporal signal of the resulting ML phylogeny was assessed using TempEst version 1.5.3 ($R^2$ = 0.26 using the best-fitting root) (50).

**Cereulide synthetase acquisition and loss events within the evolutionary history of Group III *Bacillus cereus sensu lato* facilitate the transition between emetic and diarrheal foodborne pathogen**

Using the ST 26 core SNP alignment as input, BEAST version 2.5.1 (51, 52) was used to construct a tip-dated phylogeny, where tip dates corresponded to year of isolation. For genomes that could be assigned an exact year of isolation (see section "Acquisition of Group III *Bacillus cereus s.l.* genomes and metadata" above), a fixed tip date was used (i.e., the tip date was not estimated). For genomes that could not be assigned an exact year of isolation, tip dates were estimated using a uniform distribution, with bounds selected corresponding to (i) a year beyond the maximum year of isolation (upper bound; see "Acquisition of *Bacillus cereus s.l.* genomes and metadata" section above), and (ii) a high-confidence minimum value based on available metadata (e.g., a publication reporting that a strain was isolated within a particular timeframe, but no exact year was reported for the isolate), or, if none was available, a value of 1900 (lower bound). Additionally, an ascertainment bias correction based on the GC content of the closed chromosome of emetic *B. cereus s.l.* ST 26 str. AH187 was used to account for the use of solely variant sites (53). A relaxed lognormal molecular clock (54) was used to account for varying evolutionary rates along lineages and the relatively low $R^2$ value produced when assessing the temporal signal. Due to a lack of prior information about the Group III *B. cereus s.l.* evolutionary rate, an initial clock rate of $1.0 \times 10^{-9}$ substitutions/site/year was used, along with a broad lognormal prior on the ucldMean parameter (in real space, M = $1.0 \times 10^{-3}$ and S = 4.0), which yielded a median of $3.35 \times 10^{-7}$ and 2.5 and 97.5% quantiles of $1.32 \times 10^{-10}$ and $8.52 \times 10^{-4}$ substitutions/site/year, respectively. For the substitution model, the Standard_TVMef model implemented in the SSM package (55) was used, as it was the optimal substitution model selected using the modelTest function in R's phangorn (56) package (based on BIC values), along with the Gamma category count set to 5. To explicitly account for potential sampling biases stemming from the overrepresentation of ST 26 strains isolated in recent years (i.e., 1972-

2016), a serial Birth-Death Skyline population model was used (57). A change point was introduced into the model so that the "samplingProportion" parameter could be set to 0 before the first sample date (to account for the fact that little-to-no sampling effort was made prior to the 1970s).

Five independent runs using the model described above were performed, using chain lengths of at least 100 million generations, sampling every 10,000 generations. For each independent replicate, Tracer version 1.7.1 (58) was used to ensure that each parameter had mixed adequately with 10% burn-in. LogCombiner-2 was used to combine log and tree files for each of the five independent runs with 10% burn-in, and Tracer was again used to ensure that the combined log file showcased adequate mixing with 10% burn-in. The prior was additionally sampled in the absence of sequence data, and the resulting parameter distributions were compared to the combined log file in Tracer. TreeAnnotator-2 (59) was used to produce a maximum clade credibility tree from the combined tree files, using Common Ancestor node heights. The resulting phylogeny was annotated using FigTree version 1.4.3 (60) and the phytools (25), ggtree (61, 62), and ape (22, 23) packages in R version 3.6.1 (24).

**Cereulide synthetase ancestral state reconstruction for ST 26 genomes.** Ancestral state reconstruction as it related to cereulide production capabilities was performed using the temporal phylogeny constructed using Snippy, BEAST 2, LogCombiner-2, and TreeAnnotator-2 (see section "Construction of Group III *B. cereus s.l.* ST 26 temporal phylogeny" above) as input. Stochastic character maps were simulated on the phylogeny using the make.simmap function, the ARD model, and one of the following three priors on the root node, corresponding to the *ces*-positive and *ces*-negative state of the root node: (i) equal probability of the root node belonging to a *ces*-positive or *ces*-negative state; (ii) estimated probabilities of the root node belonging to a *ces*-positive or *ces*-negative state, obtained using the make.simmap function; and (iii) probability of the root node being in a *ces*-positive or *ces*-negative state set to 0.2 and 0.8, respectively, as the probability of the ST 26 ancestor being *ces*-negative was estimated to be between 0.78 and 0.82 (depending on the choice of outgroup) when core SNPs among all Group III *B. cereus s.l.* genomes were used for ancestral state reconstruction (see section "Construction of Group III *B. cereus s.l.* maximum likelihood phylogenies and ancestral state reconstruction" above). An empirical Bayes approach was used, in which a continuous-time reversible Markov model was fitted, followed by 10,000 simulations of stochastic character histories using the fitted model and the tree tip states. The resulting phylogenies were plotted using the densityMap function in the phytools package.

**Evaluation of the influence of reference genome selection on ST 26 phylogenomic topology.** To determine if choice of reference genome affected the topology of the ST 26 phylogeny, SNPs were identified among all 64 Group III *B. cereus s.l.* genomes which belonged to ST 26 using four different reference-based SNP calling pipelines, chosen for their ability to utilize assembled genomes or both assembled genomes and Illumina reads as input: (i) BactSNP version 1.1.0 (63), (ii) Lyve-SET version 1.1.4g (64), (iii) Parsnp version 1.2 (65), and (iv) Snippy version 4.3.6. For the BactSNP, Lyve-SET, and Snippy pipelines, which can utilize both Illumina reads and assembled genomes as input, Illumina reads were used for those isolates for which they were available ($n$ = 32; Trimmomatic and FastQC were used for preprocessing, as described in section "Construction of Group III *B. cereus s.l.* ST 26 temporal phylogeny" above), and assembled genomes were used for the remaining isolates (*n* = 32). For Parsnp, which relies on assembled genomes as input, all 64 ST 26 genome assemblies were used as input.

For the BactSNP pipeline, all default steps were run as outlined in the manual. Gubbins was used to remove recombination events within the resulting pseudogenome alignment (pseudo_genomes_wo_ref.fa), and snp-sites was used to obtain an alignment of SNPs. For the Lyve-SET pipeline, all default steps were run as outlined in the manual. The resulting SNP alignment (out.informative.fasta) was queried using snp-sites to obtain an alignment of core SNPs. For the Snippy pipeline, steps were run as outlined above (see section "Construction of Group III *B. cereus s.l.* ST 26 temporal phylogeny"), with Gubbins and snp-sites used to create a core SNP alignment. For the Parsnp pipeline, core SNPs were identified using assembled genomes as input, and Parsnp's implementation of PhiPack (66) was used to remove recombination events. For each SNP alignment identified with each pipeline, IQ-TREE was used to construct a ML phylogeny using the optimal ascertainment bias-aware nucleotide substitution model selected using ModelFinder and 1,000 replicates of the ultrafast bootstrap approximation. The dist.gene function in the ape package was used to calculate the number of pairwise SNP differences between each genome in each alignment.

Each of the four reference-based SNP calling pipelines described above was run six separate times, each time using one of the following Group III *B. cereus s.l.* genomes as a reference: (i) the complete, closed chromosome of emetic *B. cereus s.l.* ST 26 str. AH187 (obtained from a human clinical isolate associated with a 1972 emetic outbreak in the United Kingdom, and previously shown to serve as an adequate reference genome for reference-based SNP calling among emetic ST 26 genomes; NCBI RefSeq Accession NC_011658.1) (9); (ii) the scaffolded draft genome of emetic *B. cereus s.l.* ST 26 str. IS195 (isolated from a pigmy shrew in Poland, and less closely related to other ST 26 isolates than *B. cereus s.l.* str. AH187; NCBI RefSeq Accession GCF_000399225.1) (67-70); (iii) the contigs of emetic *B. cereus s.l.* ST 144 str. MB.17 (isolated from food in Munich, Germany; NCBI RefSeq Accession GCF_001566445.1) (71); (iv) the contigs of emetic *B. cereus s.l.* ST 2056 str. MB.18 (isolated from food in Munich, Germany; NCBI RefSeq Accession GCF_001566385.1) (71); (v) the contigs of emetic *B. cereus s.l.* ST 869 str. MB.22 (isolated from food in Munich, Germany; NCBI RefSeq Accession GCF_001566535.1) (71); (vi) the scaffolded draft genome of emetic *B. cereus s.l.* ST 164 str. AND1407 (isolated from black currants in Denmark; NCBI RefSeq Accession GCF_000290995.1) (72, 73). This set of tested reference genomes represented all Group III STs in which cereulide synthetase-encoding genes were detected.

For each of the four SNP calling pipelines, the phylogeny constructed using SNPs identified with emetic *B. cereus s.l.* ST 26 str. AH187 as a reference genome was treated as a reference tree. The Kendall-Colijn (74, 75) test described by Katz et al. (64) was used to compare the topology of each tree constructed with SNPs identified using each of the remaining five reference genomes (emetic Group III *B. cereus s.l.* strains IS195, MB.17, MB.18, MB.22, and AND1407, representing STs 26, 144, 2056, 869, and 164, respectively) to the AH187 reference phylogeny. For each query-reference tree combination, the Kendall-Colijn test was performed using midpoint-rooted trees, a lambda value of 0 (to give weight to tree topology, rather than branch lengths), a background distribution of 100,000 random trees (64), and the following R packages: treespace (76), phangorn, ggplot2 (77), stringr (78), docopt (79), ips (80).

10. Jain C, Rodriguez RL, Phillippy AM, Konstantinidis KT, Aluru S. 2018. High throughput ANI analysis of 90K prokaryotic genomes reveals clear species boundaries. Nat Commun 9:5114.

11. Barrett T, Clark K, Gevorgyan R, Gorelenkov V, Gribov E, Karsch-Mizrachi I, Kimelman M, Pruitt KD, Resenchuk S, Tatusova T, Yaschenko E, Ostell J. 2012. BioProject and BioSample databases at NCBI: facilitating capture and organization of metadata. Nucleic Acids Res 40:D57-63.

12. Guinebretiere MH, Velge P, Couvert O, Carlin F, Debuyser ML, Nguyen-The C. 2010. Ability of *Bacillus cereus* group strains to cause food poisoning varies according to phylogenetic affiliation (groups I to VII) rather than species affiliation. J Clin Microbiol 48:3388-91.

13. Jolley KA, Maiden MC. 2010. BIGSdb: Scalable analysis of bacterial genome variation at the population level. BMC Bioinformatics 11:595.

14. Vangay P, Fugett EB, Sun Q, Wiedmann M. 2013. Food microbe tracker: a web-based tool for storage and comparison of food-associated microbes. J Food Prot 76:283-94.

15. Gardner SN, Hall BG. 2013. When whole-genome alignments just won't work: kSNP v2 software for alignment-free SNP discovery and phylogenetics of hundreds of microbial genomes. PLoS One 8:e81760.

16. Gardner SN, Slezak T, Hall BG. 2015. kSNP3.0: SNP detection and phylogenetic analysis of genomes without genome alignment or reference genome. Bioinformatics 31:2877-8.

17. Nguyen LT, Schmidt HA, von Haeseler A, Minh BQ. 2015. IQ-TREE: a fast and effective stochastic algorithm for estimating maximum-likelihood phylogenies. Mol Biol Evol 32:268-74.

18. Lewis PO. 2001. A likelihood approach to estimating phylogeny from discrete morphological character data. Syst Biol 50:913-25.

19. Kalyaanamoorthy S, Minh BQ, Wong TKF, von Haeseler A, Jermiin LS. 2017. ModelFinder: fast model selection for accurate phylogenetic estimates. Nat Methods 14:587-589.

20. Minh BQ, Nguyen MA, von Haeseler A. 2013. Ultrafast approximation for phylogenetic bootstrap. Mol Biol Evol 30:1188-95.

21. Hoang DT, Chernomor O, von Haeseler A, Minh BQ, Vinh LS. 2018. UFBoot2: Improving the Ultrafast Bootstrap Approximation. Mol Biol Evol 35:518-522.

22. Paradis E, Claude J, Strimmer K. 2004. APE: Analyses of Phylogenetics and Evolution in R language. Bioinformatics 20:289-90.

23. Paradis E, Schliep K. 2019. ape 5.0: an environment for modern phylogenetics and evolutionary analyses in R. Bioinformatics 35:526-528.

24. R Core Team. 2019. R: A Language and Environment for Statistical Computing, v3.6.1. R Foundation for Statistical Computing, Vienna, Austria. https://www.R-project.org/.

25. Revell LJ. 2012. phytools: an R package for phylogenetic comparative biology (and other things). Methods in Ecology and Evolution 3:217-223.

26. Tonkin-Hill G, Lees JA, Bentley SD, Frost SDW, Corander J. 2018. RhierBAPS: An R implementation of the population clustering algorithm hierBAPS. Wellcome Open Res 3:93.

27. Arevalo P, VanInsberghe D, Elsherbini J, Gore J, Polz MF. 2019. A Reverse Ecology Approach Based on a Biological Definition of Microbial Populations. Cell 178:820-834 e14.

28. Angiuoli SV, Salzberg SL. 2011. Mugsy: fast multiple alignment of closely related whole genomes. Bioinformatics 27:334-42.

29. Guindon S, Dufayard JF, Lefort V, Anisimova M, Hordijk W, Gascuel O. 2010. New algorithms and methods to estimate maximum-likelihood phylogenies: assessing the performance of PhyML 3.0. Syst Biol 59:307-21.

30. Steinegger M, Soding J. 2017. MMseqs2 enables sensitive protein sequence searching for the analysis of massive data sets. Nat Biotechnol 35:1026-1028.

31. Edgar RC. 2004. MUSCLE: multiple sequence alignment with high accuracy and high throughput. Nucleic Acids Res 32:1792-7.

32. Seemann T. 2019. Snippy: Rapid haploid variant calling and core genome alignment, v4.3.6. https://github.com/tseemann/snippy.

33. Li H, Durbin R. 2009. Fast and accurate short read alignment with Burrows-Wheeler transform. Bioinformatics 25:1754-60.

34. Li H. 2013. Aligning sequence reads, clone sequences and assembly contigs with BWA-MEM. arXiv:1303.3997.

35. Li H. 2018. Minimap2: pairwise alignment for nucleotide sequences. Bioinformatics 34:3094-3100.

36. Li H, Handsaker B, Wysoker A, Fennell T, Ruan J, Homer N, Marth G, Abecasis G, Durbin R, Genome Project Data Processing S. 2009. The Sequence Alignment/Map format and SAMtools. Bioinformatics 25:2078-9.

37. Quinlan AR. 2014. BEDTools: The Swiss-Army Tool for Genome Feature Analysis. Curr Protoc Bioinformatics 47:11 12 1-34.

38. Quinlan AR, Hall IM. 2010. BEDTools: a flexible suite of utilities for comparing genomic features. Bioinformatics 26:841-2.

39. Li H. 2011. A statistical framework for SNP calling, mutation discovery, association mapping and population genetical parameter estimation from sequencing data. Bioinformatics 27:2987-93.

40. Garrison E, Marth G. 2012. Haplotype-based variant detection from short-read sequencing. arXiv:1207.3907.

41. Cleary JG, Braithwaite R, Gaastra K, Hilbush BS, Inglis S, Irvine SA, Jackson A, Littin R, Rathod M, Ware D, Zook JM, Trigg L, De La Vega FM. 2015. Comparing Variant Call Files for Performance Benchmarking of Next-Generation Sequencing Variant Calling Pipelines. bioRxiv doi:10.1101/023754:023754.

42. Tan A, Abecasis GR, Kang HM. 2015. Unified representation of genetic variants. Bioinformatics 31:2202-4.

43. Cingolani P, Platts A, Wang le L, Coon M, Nguyen T, Wang L, Land SJ, Lu X, Ruden DM. 2012. A program for annotating and predicting the effects of single nucleotide polymorphisms, SnpEff: SNPs in the genome of *Drosophila melanogaster* strain w1118; iso-2; iso-3. Fly (Austin) 6:80-92.

44. Seemann T. 2019. samclip: Filter SAM file for soft and hard clipped alignments, v0.2. https://github.com/tseemann/samclip.

45. Li H. 2019. Seqtk: a fast and lightweight tool for processing sequences in the FASTA or FASTQ format, v1.2-r102-dirty https://github.com/lh3/seqtk.

46. Page AJ, Taylor B, Delaney AJ, Soares J, Seemann T, Keane JA, Harris SR. 2016. SNP-sites: rapid efficient extraction of SNPs from multi-FASTA alignments. Microb Genom 2:e000056.

47. Bolger AM, Lohse M, Usadel B. 2014. Trimmomatic: a flexible trimmer for Illumina sequence data. Bioinformatics 30:2114-20.

48. Andrews S. 2019. FastQC: a quality control tool for high throughput sequence data, v0.11.8. https://www.bioinformatics.babraham.ac.uk/projects/fastqc/.

49. Croucher NJ, Page AJ, Connor TR, Delaney AJ, Keane JA, Bentley SD, Parkhill J, Harris SR. 2015. Rapid phylogenetic analysis of large samples of recombinant bacterial whole genome sequences using Gubbins. Nucleic Acids Res 43:e15.

50. Rambaut A, Lam TT, Max Carvalho L, Pybus OG. 2016. Exploring the temporal structure of heterochronous sequences using TempEst (formerly Path-O-Gen). Virus Evol 2:vew007.

51. Bouckaert R, Vaughan TG, Barido-Sottani J, Duchene S, Fourment M, Gavryushkina A, Heled J, Jones G, Kuhnert D, De Maio N, Matschiner M, Mendes FK, Muller NF, Ogilvie HA, du Plessis L, Popinga A, Rambaut A, Rasmussen D, Siveroni I, Suchard MA, Wu CH, Xie D, Zhang C, Stadler T, Drummond AJ. 2019. BEAST 2.5: An advanced software platform for Bayesian evolutionary analysis. PLoS Comput Biol 15:e1006650.

52. Bouckaert R, Heled J, Kuhnert D, Vaughan T, Wu CH, Xie D, Suchard MA, Rambaut A, Drummond AJ. 2014. BEAST 2: a software platform for Bayesian evolutionary analysis. PLoS Comput Biol 10:e1003537.

53. Bouckaert R. 2014. Correcting for constant sites in BEAST2. https://groups.google.com/forum/#!topic/beast-users/QfBHMOqImFE. Accessed May 12, 2020.

54. Drummond AJ, Ho SY, Phillips MJ, Rambaut A. 2006. Relaxed phylogenetics and dating with confidence. PLoS Biol 4:e88.

55. Bouckaert R, Xie D. 2017. SSN: Standard Nucleotide Substitution Models, http://doi.org/10.5281/zenodo.995740.

56. Schliep KP. 2011. phangorn: phylogenetic analysis in R. Bioinformatics 27:592-3.

57. Stadler T, Kuhnert D, Bonhoeffer S, Drummond AJ. 2013. Birth-death skyline plot reveals temporal changes of epidemic spread in HIV and hepatitis C virus (HCV). Proc Natl Acad Sci U S A 110:228-33.

58. Rambaut A, Drummond AJ, Xie D, Baele G, Suchard MA. 2018. Posterior Summarization in Bayesian Phylogenetics Using Tracer 1.7. Syst Biol 67:901-904.

59. Heled J, Bouckaert RR. 2013. Looking for trees in the forest: summary tree from posterior samples. BMC Evol Biol 13:221.

60. Rambaut A. 2016. FigTree: a graphical viewer of phylogenetic trees, v1.4.3. http://tree.bio.ed.ac.uk/software/figtree/.

61. Yu G, Smith DK, Zhu H, Guan Y, Lam TT-Y. 2017. ggtree: an r package for visualization and annotation of phylogenetic trees with their covariates and other associated data. Methods in Ecology and Evolution 8:28-36.

62. Yu G, Lam TT, Zhu H, Guan Y. 2018. Two Methods for Mapping and Visualizing Associated Data on Phylogeny Using Ggtree. Mol Biol Evol 35:3041-3043.

63. Yoshimura D, Kajitani R, Gotoh Y, Katahira K, Okuno M, Ogura Y, Hayashi T, Itoh T. 2019. Evaluation of SNP calling methods for closely related bacterial isolates and a novel high-accuracy pipeline: BactSNP. Microb Genom 5.

64. Katz LS, Griswold T, Williams-Newkirk AJ, Wagner D, Petkau A, Sieffert C, Van Domselaar G, Deng X, Carleton HA. 2017. A Comparative Analysis of the Lyve-SET Phylogenomics Pipeline for Genomic Epidemiology of Foodborne Pathogens. Front Microbiol 8:375.

65. Treangen TJ, Ondov BD, Koren S, Phillippy AM. 2014. The Harvest suite for rapid core-genome alignment and visualization of thousands of intraspecific microbial genomes. Genome Biol 15:524.

66. Bruen TC, Philippe H, Bryant D. 2006. A simple and robust statistical test for detecting the presence of recombination. Genetics 172:2665-81.

67. Castiaux V, N'Guessan E, Swiecicka I, Delbrassinne L, Dierick K, Mahillon J. 2014. Diversity of pulsed-field gel electrophoresis patterns of cereulide-producing isolates of *Bacillus cereus* and *Bacillus weihenstephanensis*. FEMS Microbiol Lett 353:124-31.

68. Van der Auwera GA, Feldgarden M, Kolter R, Mahillon J. 2013. Whole-Genome Sequences of 94 Environmental Isolates of *Bacillus cereus Sensu Lato*. Genome Announc 1.

69. Swiecicka I, Fiedoruk K, Bednarz G. 2002. The occurrence and properties of *Bacillus thuringiensis* isolated from free-living animals. Lett Appl Microbiol 34:194-8.

70. Swiecicka I, De Vos P. 2003. Properties of *Bacillus thuringiensis* isolated from bank voles. J Appl Microbiol 94:60-4.

71. Crovadore J, Calmin G, Tonacini J, Chablais R, Schnyder B, Messelhausser U, Lefort F. 2016. Whole-Genome Sequences of Seven Strains of *Bacillus cereus* Isolated from Foodstuff or Poisoning Incidents. Genome Announc 4.

72. Hoton FM, Fornelos N, N'Guessan E, Hu X, Swiecicka I, Dierick K, Jaaskelainen E, Salkinoja-Salonen M, Mahillon J. 2009. Family portrait of *Bacillus cereus* and *Bacillus weihenstephanensis* cereulide-producing strains. Environ Microbiol Rep 1:177-83.

73. Biodefense and Emerging Infections (BEI) Research Resources Repository. 2019. *Bacillus cereus* Strain AND1407, NR-22159. https://www.beiresources.org/Catalog/Bacteria/NR-22159.aspx. Accessed December 24, 2019.

74. Kendall M, Colijn C. 2016. Mapping Phylogenetic Trees to Reveal Distinct Patterns of Evolution. Molecular Biology and Evolution 33:2735-2743.

75. Kendall M, Colijn C. 2015. A tree metric using structure and length to capture distinct phylogenetic signals. arXiv:1507.05211.

76. Jombart T, Kendall M, Almagro-Garcia J, Colijn C. 2017. treespace: Statistical exploration of landscapes of phylogenetic trees. Mol Ecol Resour 17:1385-1392.

77. Wickham H. 2016. ggplot2: Elegant Graphics for Data Analysis. Springer-Verlag New York.

78. Wickham H. 2019. stringr: Simple, Consistent Wrappers for Common String Operations, https://CRAN.R-project.org/package=stringr.

79. de Jonge E. 2018. docopt: Command-Line Interface Specification Language, https://CRAN.R-project.org/package=docopt.

80. Heibl C. 2008. PHYLOCH: R language tree plotting tools and interfaces to diverse phylogenetic software packages, http://www.christophheibl.de/Rpackages.html.
